# Comprehensive benchmarking of methods for mutation calling in circulating tumor DNA

**DOI:** 10.1101/2025.02.16.638482

**Authors:** Hanaé Carrié, Ngak Leng Sim, Pui Mun Wong, Anna Gan, Yi Ting Lau, Polly Poon, Saranya Thangaraju, Iain Tan, Yoon Sim Yap, Kiran Krishnamachari, Limsoon Wong, Anders Skanderup

**Affiliations:** Genome Institute of Singapore, Agency for Science Technology and Research; Institute of Data Science, National University of Singapore; Department of Computer Science, National University of Singapore; National Cancer Center Singapore, Singapore

## Abstract

Detection of somatic mutations in cell-free DNA (cfDNA) is challenging due to low variant allele frequencies and pronounced DNA degradation. Here, we present a novel approach and resource for benchmarking of somatic variant calling algorithms in cfDNA samples from cancer patients. Using longitudinally collected cfDNA samples from colorectal and breast cancer patients, we identify patient-matched samples with high and ultra-low circulating tumor DNA (ctDNA) levels. These sample pairs, preserving patient-specific germline and somatic haematopoiesis variant backgrounds, were used to generate dilution series capturing characteristics of bona-fide cfDNA samples. To benchmark the accuracy and limit of detection of 9 somatic variant calling algorithms, we used deep Whole Genome Sequencing (WGS, 150x) and ultra-deep Whole Exome Sequencing (WES, 2,000x) to construct a reference set of ∼37,000 Single Nucleotide Variants and ∼58,000 Insertions/Deletions. We tested methods under variable ctDNA levels and depth of sequencing, generating guidelines for method choice depending on use case. Using a machine learning approach, we further evaluated the potential of fine-tuning individual variant callers, revealing features that may improve accuracy in cfDNA samples. Overall, we present a new resource for benchmarking of somatic variant calling methods in cfDNA, providing insights on method choice to realize the potential of liquid biopsies in precision oncology.

## Introduction

Analysis of cell-free DNA (cfDNA) using high-throughput Next-Generation Sequencing (NGS) has emerged as a valuable approach in precision oncology [1]. cfDNA comprises DNA fragments that are released into the blood plasma through cell death or active secretion [2, 3]. In a cancer patient, a fraction of cfDNA, known as circulating tumour DNA (ctDNA), originates from tumor cells [4, 5], offering a minimally-invasive approach to access actionable tumor information with a simple blood draw [6]. This strategy has demonstrated potential in monitoring treatment response [7], detecting minimal residual disease [8], molecularly stratifying patients for targeted treatments [9], and identifying therapy-acquired resistance [10]. Moreover, recent methods have demonstrated how the cumulated signal from hundreds to thousands of somatic mutations identified with broad Whole Exome (WES) or Whole Genome (WGS) Sequencing can increase cancer detection sensitivity [11–15]. Key to such clinical applications is the ability to accurately identify cancer somatic mutations, specifically Single Nucleotide Variants (SNVs) and short Insertions/Deletions (INDELs), from blood plasma [16, 17]. However, most existing methods for somatic variant calling have not been designed for or benchmarked on plasma samples, leaving method choice challenging for end users.

Several factors contribute to higher noise levels in plasma cfDNA as compared to genomic DNA (gDNA) from tissue biopsies. One key factor is the low fraction of ctDNA present in most plasma samples, resulting in cancer mutations being present at very low variant allele frequencies (VAFs). cfDNA also exhibit higher levels of degradation as well as non-uniform depth of coverage due to nucleosome wrapping [18]. Other biological factors further complicate mutation detection, such as Clonal Hematopoiesis of Indeterminate Potential (CHIP) [19, 20] and potential for subclonal variation in multi-focal disease. Variant callers designed for tumor tissue samples typically do not account for these factors. While new methods have proposed novel strategies tailored to cfDNA mutation calling [21–23], the performance of these methods have not been independently tested and compared to existing variant callers designed for tumor tissue.

Previous studies [24–26] have benchmarked variant callers designed for tumor tissue samples using variant reference datasets generated for synthetic [27] and real [28] tumors. However, such large-scale reference datasets are currently lacking for plasma samples. Unfortunately, existing strategies used in tumor-tissue benchmarks are not directly applicable in the setting of cfDNA. For example, tumor-tissue benchmarks may utilize dilution series generated from mixing of tumor DNA with matched-normal DNA samples [24–26]. This strategy cannot be adopted for cfDNA since these samples have different error and fragmentation characteristics as compared to their matched normal samples comprising gDNA from white-blood cells. Recently, the FDA-led Sequencing Quality Control Phase 2 (SEQC2) project conducted a benchmark of cfDNA assays and somatic variant calls from four commercial providers [29–31]. However, this benchmark study focused on narrow targeted sequencing panels and plasma-only mutation calling, using nuclease-degraded DNA from cell lines as a surrogate for cfDNA. Thus, there is a need for novel approaches to generate large-scale and genome-wide reference datasets for variant calling using bona-fide cfDNA samples.

In this study, we addressed these research gaps to develop a comprehensive benchmarking framework for comparison of 9 somatic mutation callers applied to cfDNA samples from cancer patients. We created a novel reference dataset using longitudinal plasma samples from breast cancer (BRCA) and colorectal cancer (CRC) patients. For each patient, we identified timepoints with high and ultra-low ctDNA levels, establishing controlled dilution series preserving cfDNA fragmentation patterns as well as patient-specific germline and somatic haematopoiesis variant backgrounds. We then used deep WGS (150x) and ultra-deep WES (2,000x) to construct a reference set of ∼37,000 Single Nucleotide Variants (SNVs) and ∼58,000 Insertions/Deletions (indels). We evaluated and compared the performance of 9 somatic variant calling methods in plasma-vs-normal mode. We systematically evaluated the impact of ctDNA levels and sequencing depth on method accuracy, revealing guidelines for method choice depending on use case. Finally, we investigated the potential for fine-tuning individual methods using machine learning, revealing features that enable improved accuracy on cfDNA data.

## Results

### A novel large-scale cfDNA somatic mutation reference dataset

Benchmarking of variant callers in tumor tissue often make use of dilution series generated through in silico mixing of sequencing data from a tumor and a matched-normal DNA sample (e.g. buffy-coat) [24–26]. However, this strategy is ill-suited for plasma samples, as cfDNA (plasma) and gDNA (matched normal) have markedly different error and fragmentation characteristics. We reasoned that a cfDNA benchmark dataset instead could adopt in silico mixtures of patient-matched plasma samples, preserving cfDNA fragmentation patterns as well as patient-specific germline and somatic haematopoiesis variant backgrounds (Fig 1a-b). With this approach, a sample with very high ctDNA levels could be used to establish ground-truth mutations with high confidence for each patient (Fig 1c-d).

**Figure 1:**
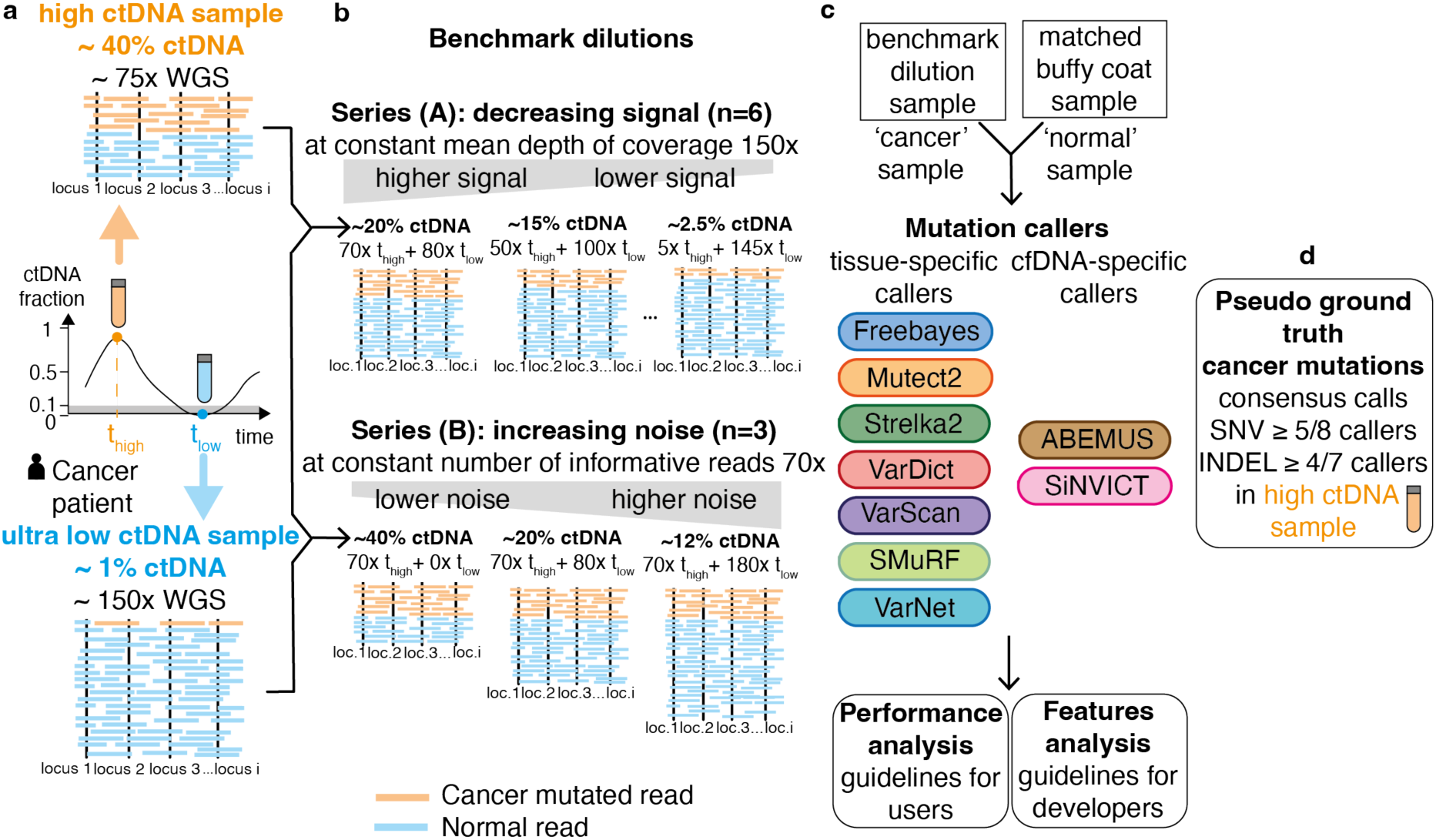
Benchmark overview. **(a)** Plasma samples with high (orange) and low (blue) ctDNA levels are identified from longitudinal samples obtained from an individual cancer patient. **(b)** Patient-specific benchmark dilutions series are created through in-silico mixing of low and high ctDNA level samples. Signal-to-noise ratios are step-wise lowered by decreasing signal (series A) or increasing noise (series B). **(c)** Nine somatic mutation callers, including seven tissue-based and two cfDNA-based methods, are applied for each benchmark dilution sample in tumor-normal mode. Performance and feature analysis are conducted to establish practical guidelines. **(d)** High-confidence cancer mutations (pseudo ground-truth) are established from a consensus over variant call methods applied to the high ctDNA level sample. ctDNA: circulating tumor DNA. cfDNA: cell-free DNA. WGS: whole genome sequencing. WBCs: white blood cells.

To create a comprehensive cfDNA benchmarking dataset based on such a strategy, we first performed deep targeted sequencing (∼5000x) and low-pass WGS (∼5x) of longitudinal plasma samples from a cohort of colorectal and breast cancer patients (Fig. 2a). In these samples, we estimated ctDNA/tumor fractions (TF) using a consensus based on copy number aberrations (ichorCNA, lpWGS) and VAFs (targeted sequencing). We identified 4 patients (1 BRCA, 3 CRC) with samples that exhibited very high (∼40%) and low ctDNA levels (∼1%) at different timepoints (Methods, Table S1). We confirmed that ctDNA levels were minimal (<3%) in all low-TF samples by inspecting VAFs for mutations identified with high confidence in the matched high-TF sample (Fig 2b).

**Figure 2:**
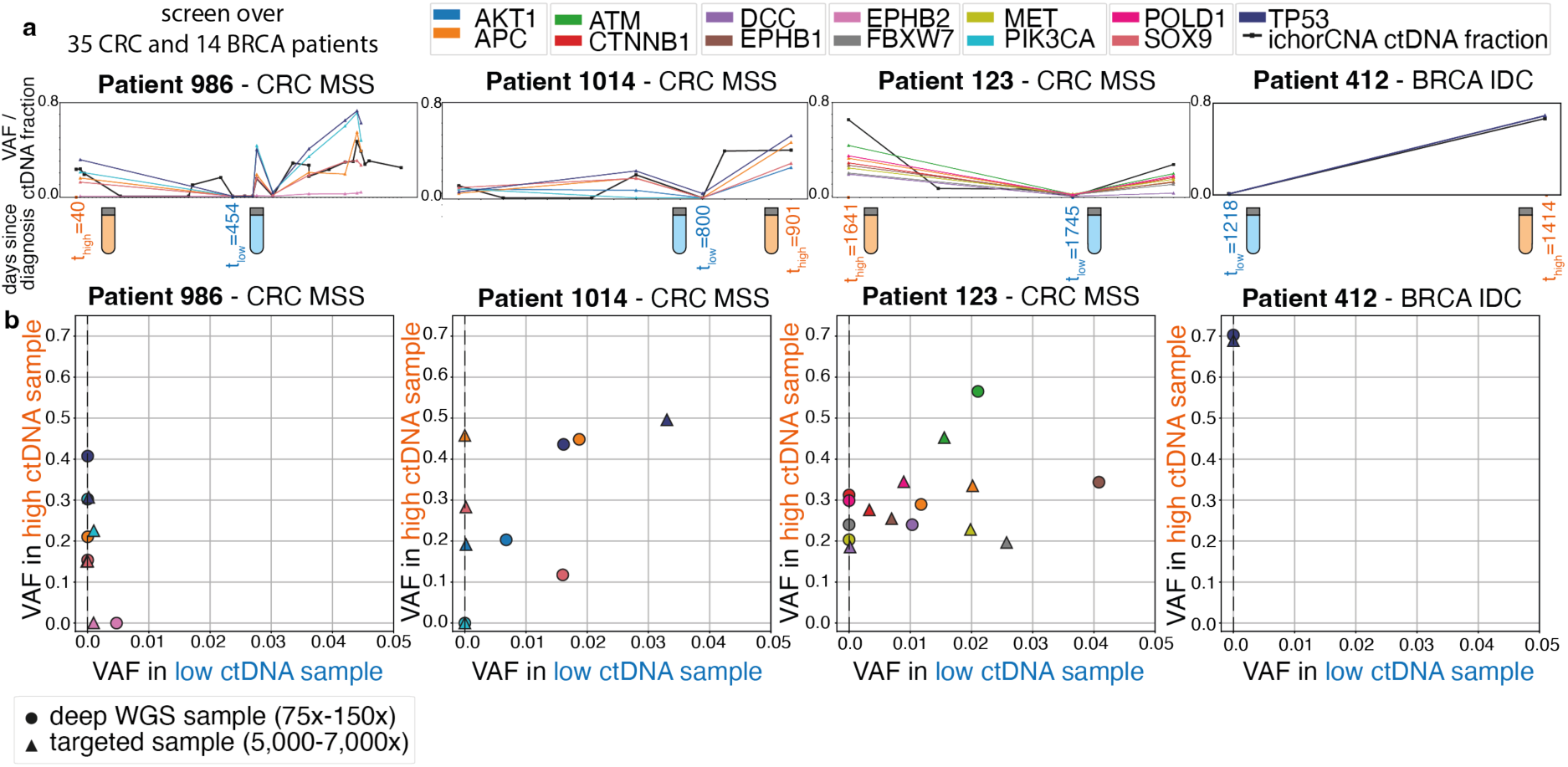
Patient selection and sample post-verification. **(a)** Sample selection: Timelines for the four candidate cancer patients (IDs = 986, 1014, 123, 412), with the timepoints for the selected high-TF sample (orange) and low-TF sample (blue) indicated. TF levels were profiled longitudinally using shallow WGS (ichorCNA) and targeted sequencing (VAFs of cancer mutations). **(b)** VAFs of cancer mutations in low and high-TF plasma samples, sequenced with both deep WGS and deep targeted sequencing. WGS: whole genome sequencing, ctDNA: circulating tumor DNA, TF: tumor fraction, VAF: variant allele frequency.

To establish dilution series, the 4 sample pairs were profiled with 150x and 75x WGS in the low and high TF sample, respectively. The low-TF sample was profiled at higher depth in order to create dilution series with lower ctDNA levels. We profiled the matched normal sample (buffy coat) for each patient using deep WGS (∼150x) to adhere to guidelines [32] advising sequencing the plasma and matched normal samples at comparable depth to limit variant calling errors due to CHIP. We created two types of WGS dilution series (Fig. 1b, Table S2): Series A was constructed with decreasing TF (i.e. decreasing signal) at fixed 150x effective depth of coverage, varying TF from ∼25% down to ∼2%. Series B was created by increasing non-ctDNA reads from the low-TF sample (i.e. increasing noise) while keeping a fixed amount of reads (70x) from the high-TF sample, thereby increasing the effective coverage from 70x to 250x in this series. In summary, both series comprise samples of high-to-low signal-to-noise ratios. However, this is achieved by decreasing signal in series A and increasing noise in series B.

### Generation of ground truths labels using consensus calling

We applied 9 SNV callers and 8 short INDEL callers to the protein-coding regions (exome) in plasma-normal mode (Fig. 1c, Table S2 and Methods). Seven of these are methods designed for tumor tissue (Freebayes [33], Mutect2 [34], Strelka2 [35], VarDict [36], Varscan2 [37], SMuRF [38], VarNet [39]) and two are methods designed for plasma cfDNA samples (ABEMUS [21], SiNVICT [23]).

We used the high-TF sample from each patient to generate high-quality ground truth variant calls using a consensus across methods (Fig. 1d, Methods). To further establish that the high-TF plasma sample could be used to define high-confidence ground-truth variants, we validated the variant calls in a patient with availability of matched tissue samples (patient 986). We compared the high-TF plasma variant calls with the corresponding tumor/normal mutation calls from deep WGS of the primary (T1) and the metastatic tissue (M1) samples. We found a highly significant overlap in these pairwise comparisons (p<10^-50^, Fisher exact test, Bonferroni correction; Fig. S1), underscoring the reliability of constructing ground truth labels using high-TF plasma samples.

In the benchmark dilution series, the VAFs of the ground-truth mutations expectedly decreased linearly as a function of dilution level (Fig. S2). Expectedly, some ground-truth variants were not detected at high dilution levels due to random sampling in the Series A dilution dataset (Fig 1b). During benchmarking, we only considered ground-truth variants supported by at least one variant read in a diluted sample, thereby omitting counting of non-sampled ground-truth variants as false-negatives.

### Variant calling accuracy at 150x sequencing depth

We considered two use cases when comparing the mutation callers: (i) unbiased genome-wide discovery, and (ii) clinical targeted genotyping. In the discovery use case, we evaluated the extent that methods offer a good trade-off between precision and recall when applied to an entire genome or exome [11–15]. In the clinical genotyping use case, the key challenge is often the ability to detect if known actionable mutations are present in a given sample [43, 44]. In this setting, we therefore evaluated the sensitivity of methods at a given fixed and relaxed precision level. In both use cases, we analysed the of impact of the two distinct dilution series with lower signal and higher noise in each of the four patients (Fig. 1b, Fig. S4).

In the discovery setting, all methods expectedly demonstrated lower accuracy when the signal-to-noise increased. Strelka2 and SMuRF exhibited the best performance for SNV calling across all ctDNA ranges, both when samples were diluted with lower signal and higher noise (Fig. 3a). The AUPRC of these two methods were about twice as high as the next-best methods across most dilution levels. The performance of individual methods were largely consistent across the four patients (Fig. 3c, Fig. S5a-b). For INDEL calling, VarScan and SMuRF consistently demonstrated highest accuracy, with Strelka2 performing well only in samples with highest signal-to-noise levels (Fig. S4). We observed consistent results when performing analysis at the whole genome level, with high concordance of method rankings between exome and whole genome calling (Fig. S7). However, the AUPRC values were almost 50% lower for all methods in the whole genome setting, highlighting the increased difficulty in calling mutations in many non-coding regions.

**Figure 3:**
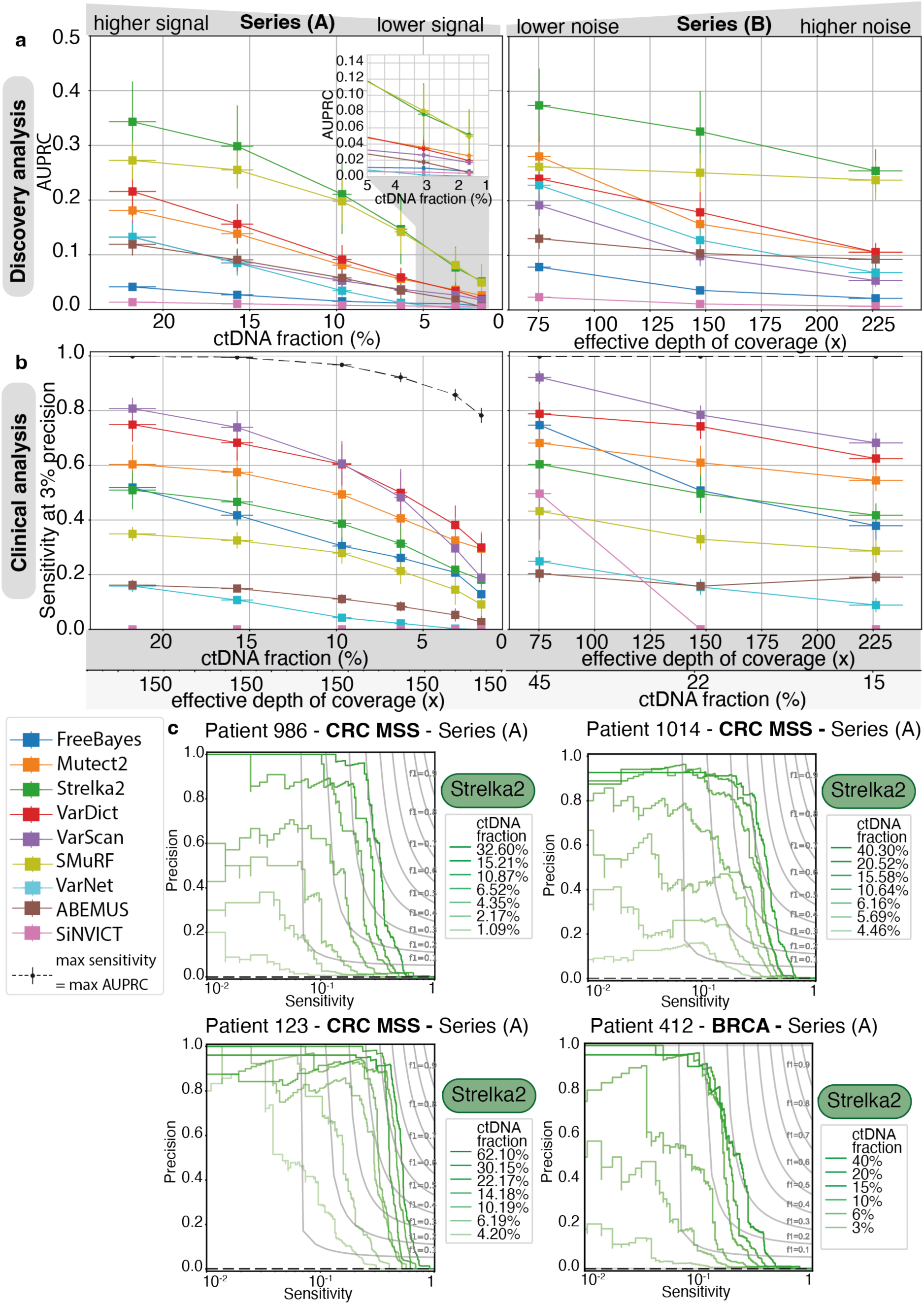
Variant calling benchmark at 150x sequencing depth. **(a,b)** Exome variant calling accuracy for each method, when applied to distinct dilution series: decreasing signal series (A), and increasing noise series (B). Methods were evaluated in two settings: **(a)** unbiased discovery analysis focusing on the AUPRC metric, and **(b)** clinical genotyping focusing on sensitivity at a fixed precision level. Vertical error bars show the standard error of the mean (s.e.m.) of the averaged metric curves, horizontal error bars show the s.e.m. of the ctDNA fraction estimates among patients (n=4). **(c)** Precision-Recall curves for Strelka2, the best performing method in the discovery setting, shown separately for each patient. The reference undiluted high ctDNA sample is also displayed as the first sample of the dilution series. Grey lines represents iso-F1 curves. SNV: Single Nucleotide Variant. TF: Tumor Fraction, ctDNA: circulating tumor DNA. AUPRC: Area Under Precision Recall Curve.

In the clinical genotyping setting, three methods (VarScan, VarDict and Mutect2) demonstrated highest sensitivity (Fig 3b). VarScan generally showed highest accuracy, except when TF dropped below 5% in series A (Fig 3b). Mutect2 and VarDict was able to outperform VarScan in that latter case. For INDELs, VarScan showed best performance of all callers across both dilution series (Fig. S4).

Surprisingly, both cfDNA-specific callers (ABEMUS and SiNVICT) included in this study exhibited poor accuracy in these benchmarks (Fig. 3a-b). Overall, variant callers developed for tumor tissue variant discovery also showed highest accuracy for variant calling in plasma samples at intermediate sequencing depth (75-200x).

### Variant calling accuracy at 2,000x sequencing depth

To evaluate the impact of increased sequencing depth on variant calling accuracy, we re-sequenced the samples from one CRC patient using ultra-deep WES at 2,000x sequencing depth (Methods, Table S3). We generated consensus ground truth calls using the high-TF ultra-deep WES sample and obtained 739 SNVs and 1,300 INDELs. In the discovery analysis setting emphasizing overall variant calling accuracy, the ranking of callers remained similar between the 150x and 2,000x sequencing. The main exception was ABEMUS whose performance was improved, and similar to Strelka2 and SMURF, in the ultra-deep sequencing setting (Fig. 4a, Fig. S8). In the clinical analysis setting, emphasizing variant calling sensitivity, we observed notable differences in method performance with ultra-deep sequencing. Here, VarScan and ABEMUS demonstrated highest sensitivity, maintaining recall above 50% across all ctDNA ranges and noise levels (Fig. 4b, Fig. S8). In contrast, VarDict and Mutect2 exhibited a drop in sensitivity at 2,000x (20-40%) compared to 150x (40-75%, Fig. 4b, Fig. S8). Overall, our findings demonstrate that somatic mutation caller accuracy depends on factors such as use case and sequencing depth.

**Figure 4:**
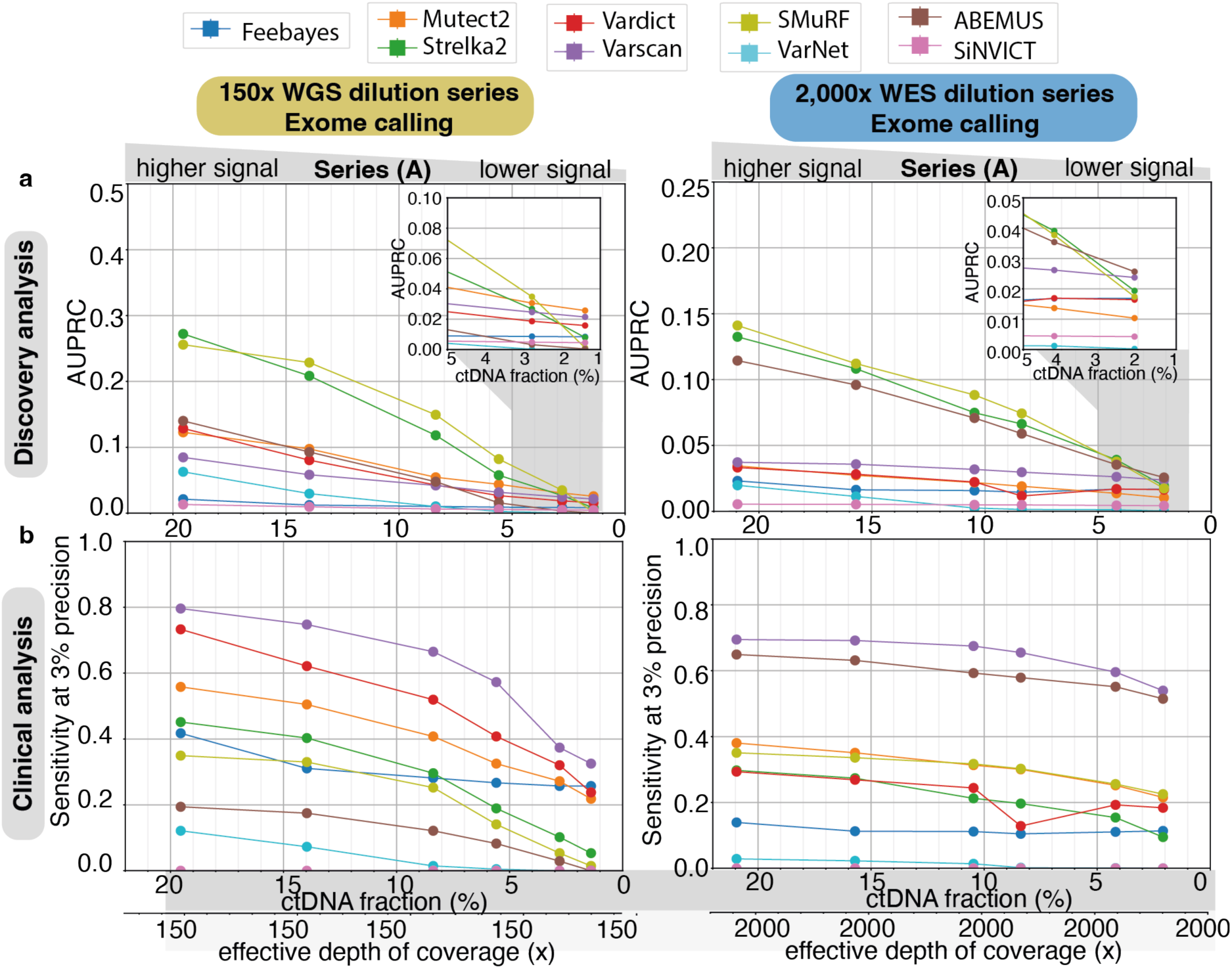
Impact of sequencing depth on variant calling accuracy. Callers’ performance on Single Nucleotide Variants in exonic regions in dilution series (A) at two different fixed coverage levels: 150x (left, derived from Whole Genome Sequencing samples) and 2,000x (right, derived from Whole Exome Sequencing samples). Callers are evaluated in two application settings: **(a)** discovery analysis with the Area Under Precision-Recall Curve metric with a zoomed version of the ctDNA fraction range [1-5%] and **(b)** clinical analysis with the sensitivity at a precision fixed to 3%. ctDNA: circulating tumor DNA.

### Fine-tuning tumor tissue-designed callers for cfDNA

Our benchmark revealed that variant callers designed for tumor tissue analysis can outperform callers specifically designed for cfDNA. We therefore investigated to what extent these tissue-based variant callers could be further fine-tuned to improve performance on cfDNA. We leveraged the generated cfDNA variant calling benchmark dataset to train cfDNA-tuned versions of each variant caller. We trained Random Forest classifiers using each caller’s predicted variant score combined with other auxiliary features provided in the Variant Call Format (VCF) output. VarNet was excluded from this analysis due to its absence of companion features due to its image-based convolutional neural network approach. We trained models on the combined 150x benchmark dataset based on the three CRC patients using leave-one-subject-out cross-validation. Notably, this approach resulted in a marked improvement in the precision of Mutect2, positioning it as the top-performing method at 150x for AUPRC, surpassing both Strelka2 and SMuRF (Fig. 5a,b). The cfDNA-tuned FreeBayes also demonstrated improvements in both AUPRC and sensitivity but remained in the mid-tier of method rankings. Strikingly, fine-tuning on cfDNA data did only yield improvements in sensitivity for the remaining methods (Fig. 5c). This underscores the need for innovative approaches to account for cfDNA-specific sequencing properties in order to further enhance sensitivity.

**Figure 5:**
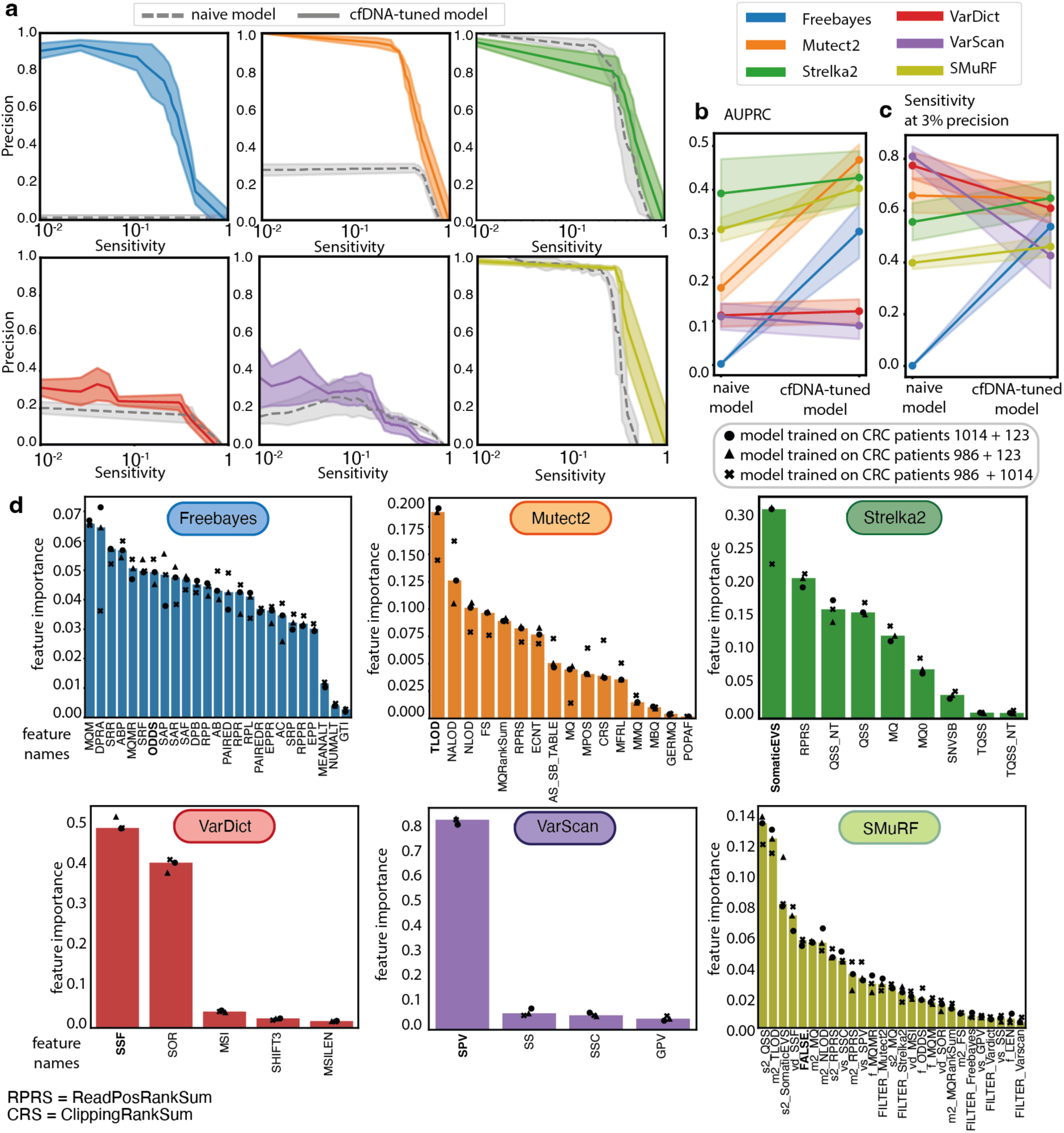
Fine-tuning variant calling methods for cell-free DNA. **(a)** We fine-tuned and evaluated the performance of tissue-based variant calling methods using a Random Forest model and leave-one-patient-out cross-validation (150x sequencing depth). The averaged Precision-Recall curves display both the performance of the initial base callers designed for tumor tissue (dashed lines) and of the corresponding cfDNA-tuned methods (coloured lines) on the unseen patient data. **(b)** Gain in AUPRC between the naive and the corresponding cfDNA-tuned caller. **(c)** Gain in sensitivity at 3% precision between the naive and the corresponding cfDNA-tuned caller. The error bands for **(a-b-c)** show the standard error of the mean (s.e.m.) of the three metrics values created by permutations of the three CRC patients. **(d)** Bar-plots display for each cfDNA-tuned caller the median features importance based on the Mean Decrease in Impurity to predict pseudo ground truth labels, points show values for cross-validation evaluation. The feature name corresponding to the prediction score of the naive model is displayed in bold. cfDNA: cell-free DNA, CRC: colorectal cancer, AUPRC: Area Under Precision-Recall Curve.

To further explore which companion features enabled performance gains on cfDNA, we ranked features according to their importance, determined as the mean of the impurity decrease within the Random Forest models fitted during cross-validation (Fig. 5d, Methods). This analysis revealed that, for all callers, except FreeBayes and SMuRF, the standard prediction score is the most important feature. However, the cfDNA-tuned version of Freebayes utilized many other features, including mapping quality, strand bias, VAF, read counts, and base quality. The cfDNA-tuned Mutect2, exhibiting significantly improved precision, incorporated multiple features derived from the matched normal sample. Notably, the NALOD feature, defined as the likelihood of an artifact to be present in buffy coat with same VAF as in plasma, may aid in filtering potential CHIP variants. The cfDNA-tuned version of Strelka2 incorporated the variant position in the read, quality scores, and mapping quality in both plasma and the matched normal. The cfDNA-tuned SMuRF relies heavily on its own standard prediction score, as well as those of Strelka2, Mutect2, and VarDict, with additional incorporation of features such as the quality score of Strelka2 and Mutect2’s mapping quality and matched normal genotype likelihood. In contrast, VarDict and VarScan did not rely on companion features in the ctDNA-tuned versions, suggesting minimal improvement based on the existing auxiliary features used by these methods.

Overall, this analysis highlights the fine-tuning potential of each method designed for tumor tissue for further improved performance for cfDNA variant calling. Inspection of feature importance yields further insights for future cfDNA-specific software development.

### Guidelines for accurate whole genome somatic variant calling in cfDNA

Our benchmark dataset allowed us to conduct an exhaustive performance analysis of tumor-agnostic cfDNA variant calling approaches. Our findings led us to derive practical guidelines for users to optimize somatic mutation calling in cfDNA (Fig. 6). Notably, our experiments showed that for unbiased mutation discovery analysis (∼100x coverage), Strelka2 and SMuRF provided the most accurate SNV calls, while VarScan yielded the most accurate INDELs calls. For clinical analysis, emphasizing sensitivity and traditionally conducted at high depth (>1,000x), VarScan and ABEMUS achieved the highest sensitivity. Furthermore, our analysis highlighted Mutect2 and VarDict in the mid-range of caller rankings across all cfDNA experiments. Moreover, Mutect2 also demonstrated superior performance in whole genome analyses at low ctDNA concentrations (Fig. S7), and showed potential for improved precision following cfDNA-specific fine-tuning (Fig. 5a-b).

**Figure 6:**
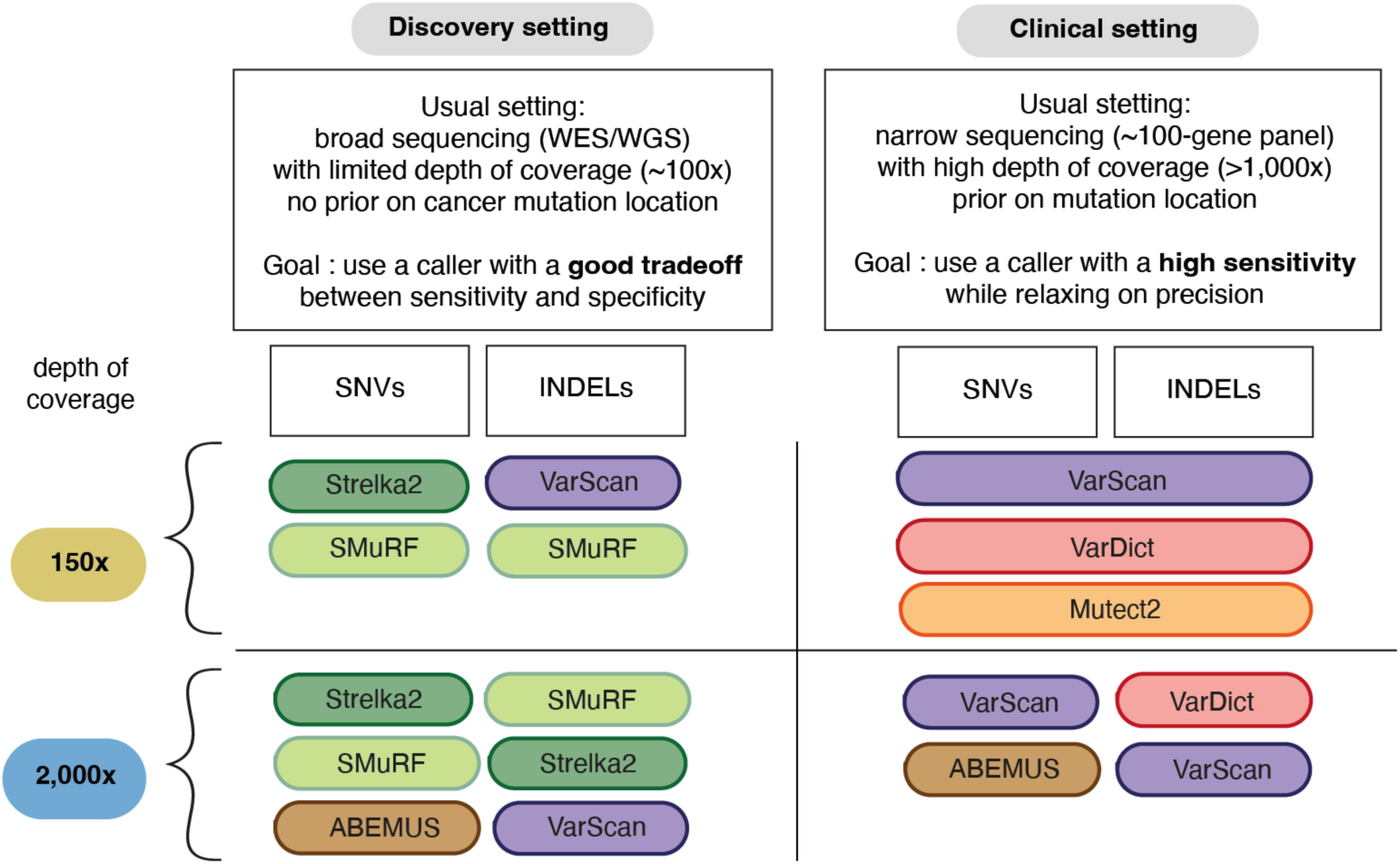
Guidelines for variant calling in cell-free DNA samples. Top-performing method depending on the analysis setting: (left) unbiased discovery analysis, (right) clinical analysis; (top) intermediate sequencing depth (∼150x), (bottom) deep sequencing (∼2,000x).

## Discussion

The emergence of cfDNA analysis using NGS has offered promising results for precision oncology, enabling non-invasive and repeatable profiling of tumor molecular information. Indeed, accurate tumor-agnostic detection of cancer somatic mutations from blood samples has important use cases such as early non-invasive genotyping of tumors, discovery and analysis of tumor evolution and treatment resistance, as well as early cancer detection. However, the absence of cfDNA benchmark datasets and unbiased evaluation of existing methods presents an unmet need to the field. Here, we tackle this challenge with three main contributions: the creation of a large-scale benchmark dataset using a novel strategy based on longitudinal patient-matched samples pairs; the evaluation and comparison of variant callers in both discovery and clinical analysis settings; evaluation of fine-tuning potential of existing methods using a machine-learning approach.

First, we presented a large-scale benchmark dataset created from real CRC and BRCA plasma samples for somatic mutation calling. Our dataset encompasses two widespread cancer types and contains in total close to 100,000 high-quality SNVs and short INDELs. This dataset is complementary to the SEQC2 studies [29, 30]. The SEQC2 studies focus on narrow targeted assays from molecular diagnostic companies, yielding in total only about 100 cancer mutations for benchmark studies. Moreover, the SEQC2 studies use mock cfDNA samples created from cell-lines [31]. To mimic cfDNA degraded fragments, the reference cell lines were mixed in vitro at known ratios and enzymatically sheared and size selected. While this approach gives the advantage of creating well-characterized and verified ground truths using orthogonal approaches, most of the ground truths are likely germline mutations rather than bona fide cancer somatic mutations. Indeed, 91% of verified mutations happened overlapped annotated germline mutations in dbSNP [50]. Our proposed dataset therefore provides a complementary large-scale dataset of bona fide cancer mutations derived from real cfDNA samples, particularly in genome/exome-wide sequencing settings with a matched normal sample.

Our analysis let led us to derive practical guidelines for users to optimize their choice of somatic mutation calling for cfDNA. Importantly, the benchmark demonstrated that distinct methods were superior in the unbiased discovery and clinical setting emphasising sensitivity. Moreover, method performance varied further when considering SNV and INDEL calling as well as depth of sequencing (150 vs 2,000x). Finally, using a machine learning approach, we evaluated the fine-tuning potential of each method for further improving performance for cfDNA variant calling. Inspection of feature importance yielded insights for future cfDNA-specific software development.

In addition to performance metrics, the practicality of implementation and hardware requirements are crucial considerations when deploying software applications. Therefore, we meticulously documented the design and implementation strengths and weaknesses of each method and accompanying software (Table S4). Methods designed for tumor tissue were in general easy to run on large-scale datasets. As for cfDNA-specific methods, ABEMUS requires a panel of at least ten normal buffy coat samples sequenced with the same platform and at the same depth in order to build the background error model. Such panels are rarely available in practice. SiNVICT does not output a probability score but several lists of calls with different filters applied, which may be inconvenient for users. We recommend that it is crucial for mutation callers to provide default parameters for users to be able to run on new samples and report a probability score for each mutation call rather than just a binary label.

The constructed dataset and benchmark methodology present certain limitations and opportunities for further improvement. First, the ultra-low ctDNA sample used to dilute the high ctDNA sample were not entirely cancer-free cfDNA. However, it remained the best proxy for a patient-matched non-cancer cfDNA sample. Despite accounting for the presence of residual ctDNA when estimating the resulting TF, there might still be slight genetic differences between the high and ultra-low ctDNA samples due to treatment or tumor evolution during the time gap. Treatment-related mutations can only emerge when the ultra-low ctDNA sample follows the high ctDNA sample. This scenario is observed in two of the four studied patients (Patients 986 and 123) with time gaps of 1 year and 3 months, and 3 months, respectively. Although this may introduce some noise in the benchmark dataset, the impact is mitigated by the ultra-low levels of ctDNA.

The newly created dataset and benchmarking strategy introduced in this study may also benefit future research in evaluating callers of other molecular alterations, such as structural variants and copy number alterations. It could also be used as a training or validation set for development of novel mutation callers. Overall, our study provides a novel framework to improve and standardise cancer mutation detection in blood, supporting the development of next-generation liquid biopsy assays and algorithms for precision oncology.

## Methods

### Patient sample preparation

Plasma and patient-matched buffy coat samples were isolated from whole blood within six hours from collection by centrifuge 300xg 10min at room temperature. Upper plasma layer was further centrifuged at 9720xg for 10min at 4°C and subsequently stored the supernatant at -80°C. Middle Buffy coat layer from first spin was removed and stored directly at -80°C until further process. Tissue was cut into pieces immediately after collection from Operation Theater and snap freeze with Liquid nitrogen before storage at -80°C.

### DNA extraction

Different kits were used for DNA extraction: Plasma(cfDNA) used Qiagen QIAamp Circulating Nucleic Acid Kit; Buffy coat (gDNA) used Qiagen QIamp DNA Blood Midi and tissue used Qiagen AllPrep DNA/RNA Mini Kit. DNA was further quantified with Invitrogen Quant-iT™ PicoGreen ® dsDNA Reagent.

### Library Preparation and NGS sequencing

Up to 100ng of tissue or Buffy coat gDNA and 4-50ng of cfDNA were used for library prep. Tissue DNA & Buffy coat gDNA was sheared to 200bp by Covaris LE220 Focused-ultrasonicator with the following condition: Target BP(Peak) 200, Duty Factor 30%, Peak Incident Power (W) 450, Cycles per Bust 200, Treatment Time 175sec. Sheared DNA subsequently was cleaned-up with 1.4X Agencourt AMPure XP beads (Beckman Coulter, A63881). Purified DNA and cfDNA (without shearing) were further processed with KAPA Hyper Prep Kit (Roche, KK8504) for libraries generation using library adapters with a random 8-mer proximal to the library index site (Oligos synthesized by IDT). Hybridization capture was done using an IDT xGen Exome (ver 1) Hyb Panel and reagents as per manufacturer’s instruction (xGen® Hybridization and Wash Kit, IDT 1080577). Libraries were quantified with KAPA Library Quantification Kits (KAPA, KK4854) and sent for 100x coverage for Whole Genome Sequencing and 1000x or 200x coverage for Whole Exome Sequencing. Sequencing was done Paired-end (2×151bp) on an illumina Novaseq 6000 S4.

### Variant calling in cfDNA deep targeted sequencing (∼5,000x) samples

The deep targeted sequencing cfDNA samples were analysed using the bcbio-nextgen pipeline [51], including read alignment with BWA mem, PCR duplicate marking with biobambam, as well as recalibration and realignment with GATK. Somatic variant calling was performed using MuTect and VarScan with default parameters, and all calls were annotated with Variant Effect Predictor. Variants were removed if they were outside coding regions. The inferred VAFs were either from one of the two callers if the variant was missed by one caller, or the mean if the variant was called by both callers. Variants from HLA-A, KMT2C and MUC17 were filtered because the majority of variants in these genes were also found by at least one caller at ≥0.005 VAF in buffy coat sequencing. The list of known cancer driver mutations of a given patient is composed of the union of mutations identified in at least one cfDNA sample.

### ctDNA tumor burden estimation in shallow WGS (∼5x) samples

We ran ichorCNA [42] with the same parameters as indicated in the authors’ Wiki page,, except for parameter *chrs* set to *c(1:22)* to analyse autosome only to reduce complexity and *includeHOMD* set to *True* to include HOMozygous Deletions as large bin size is used (1Mb), consistent with parameter usage description in the documentation.

### Selecting candidate cfDNA samples of patients with an exceptional timeline

Building the benchmark cfDNA dataset required precisely identifying relevant patients to send to a costly deep WGS procedure and post-verification of those samples. To identify suitable patients, we performed shallow WGS (∼5x cov) on multiple timepoints. This allowed us to estimate tumor fraction (TF) using the ichorCNA [42] method, describe in the previous paragraph. Additionally, ultra-deep targeted sequencing (∼5,000x cov) was conducted to track the VAFs of known CRC driver mutations, which were identified in at least one timepoint, as described above. To classify a sample as ‘high ctDNA’, both the ichorCNA [42] TF estimate and the VAFs of known CRC mutations needed to exceed 20%, as illustrated in Fig. 2a. We compared the ichorCNA [42] TF estimate with four purity estimation methods designed for tumor tissue samples, adapted for high TF values, to ensure the obtained estimates were consistent. For a sample to be categorized as ‘ultra-low ctDNA’, the ichorCNA [42] TF estimate has to be 0%, indicating the TF is below 3% since ichorCNA [42] detection limit is TF=3%. Additionally, the VAFs of known mutations had to range from less than 1% to up to 3%, as depicted in Fig. 2a.

### ctDNA tumor burden estimation in deep WGS (∼100x) samples

We first ran on all initial deep WGS cfDNA samples the cfDNA-specific ichorCNA [42] TF estimator. We also reported for each initial sample the median VAF of known cancer driver mutations (Table S1). For the high ctDNA samples, we also ran the four tumor purity estimators designed for tissue samples: THetA2 [52], TitanCNA [53], AbsCN-seq [54] and PurBayes [55]. As the amount of tumor-derived reads in those high ctDNA samples is comparable to what is observed in a tissue sample, they are expected to give values comparable to ichorCNA [42] TF. Those tumor purity estimators were applied in tumor-normal mode on cfDNA and using somatic mutations and copy number alterations called by SMuRF [38] and CNVkit [56], respectively, via the bcbio-nextgen [51] workflow. All high ctDNA samples included in the study had comparable tissue and plasma-based ctDNA fractions. The TF of the high ctDNA samples was set as the mean between ichorCNA [42] TF and the median of the four purity estimates. For ultra-low TF samples, as we observed a discrepancy between the ichorCNA [42] TF estimated on matched deep VS shallow WGS samples, we relied on the mutations’ VAF to estimate TF. The TF of the ultra-low ctDNA samples was set as TF = 2 x median_VAF_. The intermediate and final values of TF estimates for each initial sample is summarized in Table S1. The study thus includes three matched pairs of high ctDNA and ultra-low ctDNA samples with the following estimated TFs. For patient 986, TF^high^ =32.6% and TF_low_ =0%. For patient 1014, TF_high_ =40.3% and TF_low_ =2.56%. For patient 123, TF_high_=62.1% and TF_low_=3.15%. For patient 412, TF_high_ ∼40%, TF_low_ ∼0%.

### Verifying deep WGS candidate cfDNA samples of patients

Once the candidate samples were sequenced with deep WGS, we verified our conditions on TF estimate and mutations’ VAFs were satisfied on the obtained sample and provided reliable TF estimates for each sample in Table S1. To verify the condition on the TF estimate, we applied the TF estimation methods on the deep WGS samples. We obtained expected elevated values for high ctDNA samples (range TF=[23-65%]) and then set the TF estimate of high ctDNA samples as the mean between ichorCNA [42] TF and the median of 4 DNA purity estimates. For ultra-low TF samples, we observed a discrepancy between the ichorCNA [42] TF estimates obtained on the deep WGS (range TF=[3-8%]) compared to the matched shallow WGS (TF=0%). Thus, we decided to rely on the mutations’ VAF to estimate TF in those ultra-low TF samples. To check the condition on mutations’ VAFs is satisfied, Fig. 2b reports the VAFs of mutations attributed to this patient obtained in the deep WGS samples. As desired, the VAFs are concentrated in the upper left corner of the scatter plot, indicating a pronounced prevalence in the high ctDNA sample and minimal incidence in the ultra-low ctDNA sample. To set a confident TF estimate for the ultra-low ctDNA samples, TF was set as 2 x median_VAF_.

### Benchmark dilution series creation

Samples were processed using the bcbio-nextgen [51] pipeline. Sequencing reads were aligned to the GRCh37 reference human genome using BWA-MEM v0.7.17. Samtools v.1.9 was used to perform *in silico* mixture series from the high and ultra-low ctDNA samples with ‘samtools view -s’ and ‘samtools merge’ commands. The command ‘samtools depth -a’ was used to estimate average depth of coverage.

### Germline mutation calling

Genome Analysis Toolkit (GATK) [57] was run as part of the bcbio-nextgen [42] pipeline to call germline mutations on the matched normal sample of each plasma sample. Those germline calls were systematically removed from analysis to focus on cancer somatic mutations.

### Somatic mutation calling methods

Each caller was applied with default recommended parameters and following the authors’ guidelines unless otherwise specified. For the five callers designed for tumor tissue samples Freebayes v1.3.5, Mutect2 v2.2, Strelka2 v2.9.10, VarDict v1.8.2 and VarScan 2.4.4, the bcbio-nextgen [51] framework v1.2.9 was used to generate somatic mutation calls. Calling was run in tumor-normal mode, without INDEL recalibration. Duplicates were marked using biobambam v2.0.87. We lowered the VAF threshold ‘min_allele_frequency’ (default 10%) to adapt it to the low VAF mutations harboured in cfDNA. Thus, the VAF cutoff was set to 1% for 150x series and 0.01% for 2,000x series in order to match the limit of detection of a single supporting variant. We further recovered in the post-processing of output VCF files the low VAF calls that were rejected just because of the low VAF. The four other callers, SMuRF, VarNet, ABEMUS and SiNVICT, were run independently. SMuRF v2.0.12 was applied using output of the 5 previous callers. VarNet v1.1.0 was run after marking and removing duplicates using Picard. ABEMUS required a panel of normal WBC samples sequenced with comparable depth of coverage and, more importantly, sequenced with the same sequencing assay and platform to build a background error model. Although authors recommended at least 10 samples, only 8 such samples from other CRC patients could be obtained from the studied cohort. For SiNVICT, we lowered the minimum required read depth parameter min-depth (default 100) for a given call to 5 in ∼100x WGS and to 50 in ∼1,000x WES. Mutation callers were run on High Performance Computing hardware provided by A*STAR Computational Resource Centre on Red Hat Enterprise Linux 8.1 (Optpa).

### Somatic Mutation Caller Evaluation Metrics

For this binary classification problem (mutation or no mutation for each considered genomic locus), the following notations are used: True Positive (TP), True Negative (TN), False Positive (FP), False Negative (FN) and the following wide-spread metrics are considered:

- 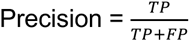
- 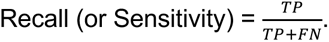
- F1−score = 2 Precision × Recall / Precision + Recall (harmonic mean of the Precision and Recall)
- Area Under the Precision Recall Curve (AUPRC) which is often use in rare-event binary classification problem. Its interpretation though is a bit harder than classical Area Under the Receiver Operating Characteristic (ROC) curve (AUC) as a random estimator AUC is 0.5 while a random estimator AUPRC depends on the proportion of positive events. For instance, if there are 1% true mutations in the dataset, the random AUPRC is 0.01.

The Precision Recall Curve (PR curve) helps visualising the performance of the individual algorithms under different variant score thresholds (Freebayes log-odds score ODDS, MuTect2 tumor log-odds score TLOD, Strelka2 tumor phred score SomaticEVS, VarDict SSF score, VarScan SSC score, 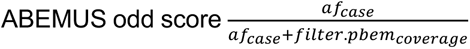). For the SiNVICT caller however, the cut-off threshold is an input parameter. The method does not output any score but reports 6 lists of calls corresponding to the successive application of 6 different filters. Thus, we assigned a score for each call equal to the number of occurrences of this given call across all lists divided by 6. Therefore, a call present even after applying all filters will be considered very confident with probability score equals to 1, while a call only present in the first list, thus caught by the Poisson model, will be considered of small confidence with a probability score equals to ⅙=0.167.

The implementation of Precision, Recall, AUPRC, PR curve provided by the scikit-learn [58] v.0.21.3 library was used in our analysis.

### Benchmark analysis and visualisation

Results analysis and visualisation was performed in Python v3.7.10. For feature analysis using Random Forest, the scikit-learn [58] v.0.21.3 library was used. The Venn diagrams were created using the nVennR package in R v4.0.4.

### Reporting summary

Further information on research design is available in the Nature Research Reporting Summary linked to this article.

## Data availability

Biological sequencing data and the created *in silico* series is available from NCBI Sequence Read Archive under a BioProject accession number, which will be provided during the publication process.

## Code availability

Script for data generation and results analysis as well as the configuration files used to run the mutation calling pipelines, will be accessible on GitHub. The repository link will be made available upon publication.

## Acknowledgements

This work was supported by the A*STAR Computational Resource Centre through the use of its high performance computing facilities.

## Author contributions

A.S., W.L.S., and H.C. designed the study. I.T., and Y.S.Y. provided samples and clinical information. P.M.W., Y.T.L., A.G., and P.P. performed the sequencing experiments. S.T. designed and ran the in-house VarDict-based mutation calling pipeline on the deep targeted cfDNA samples. H.C. created the *in silico* benchmark dilution series and built the pseudo ground truth labels. H.C. performed mutation calling on deep WES/WGS with assistance from N.L.S.. H.C performed data analysis and prepared the figures. A.S. and H.C. wrote the manuscript, with contributions from all authors.

## Supplementary Information

**Supplementary Figure 1:**
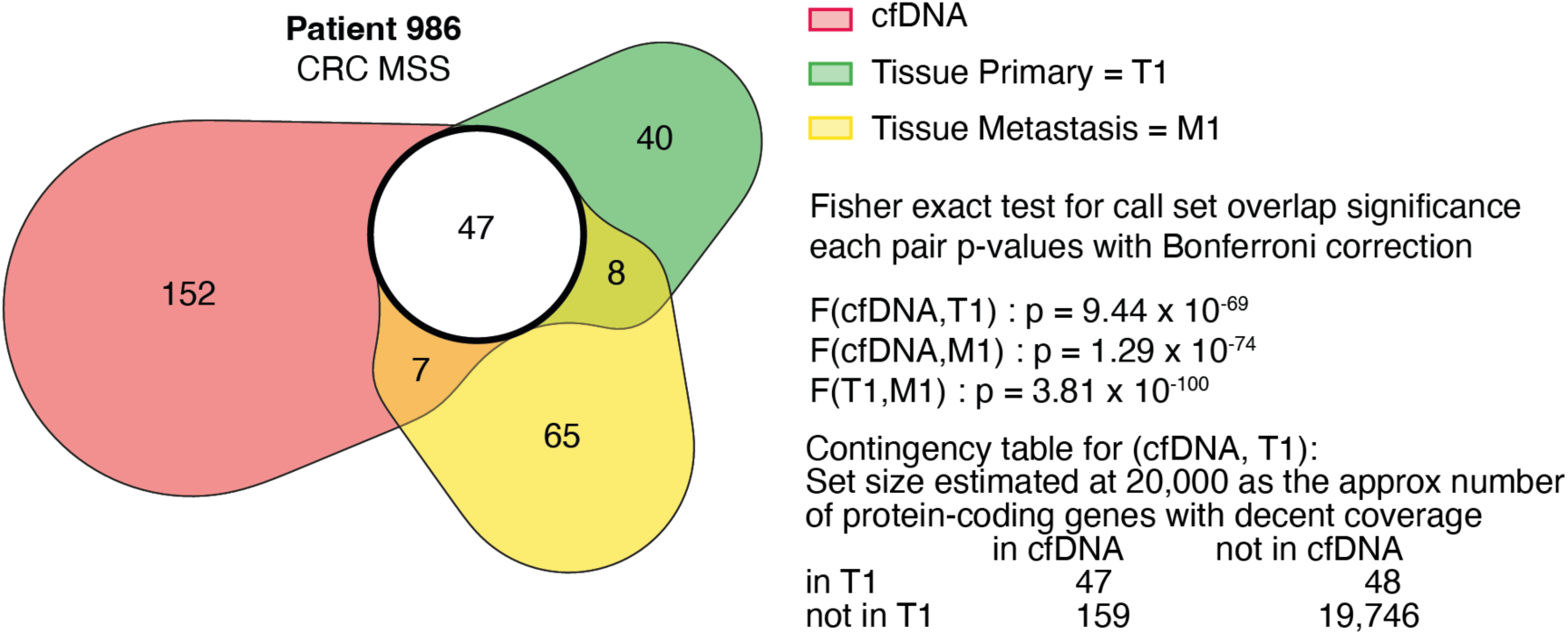
Pseudo ground truth concordance between cell-free DNA (100x, WGS) and tumor tissue (100x, WGS), restricted to the exome. Venn diagram showing the overlap in consensus exomic calls using the high ctDNA sample (K = 5 callers out of 8) vs. the tumor - primary and/or metastasis - tissue samples (K = 3 callers out of 6) for colorectal cancer patient 986. SMuRF is excluded from ground truth generation as it is an ensemble caller. For tissue samples, cfDNA-specific methods ABEMUS and SiNVICT are not applied. Fisher exact test based on contingency tables is applied on each pair of call sets to quantify overlap significance. ctDNA: circulating tumor DNA.

**Supplementary Figure 2:**
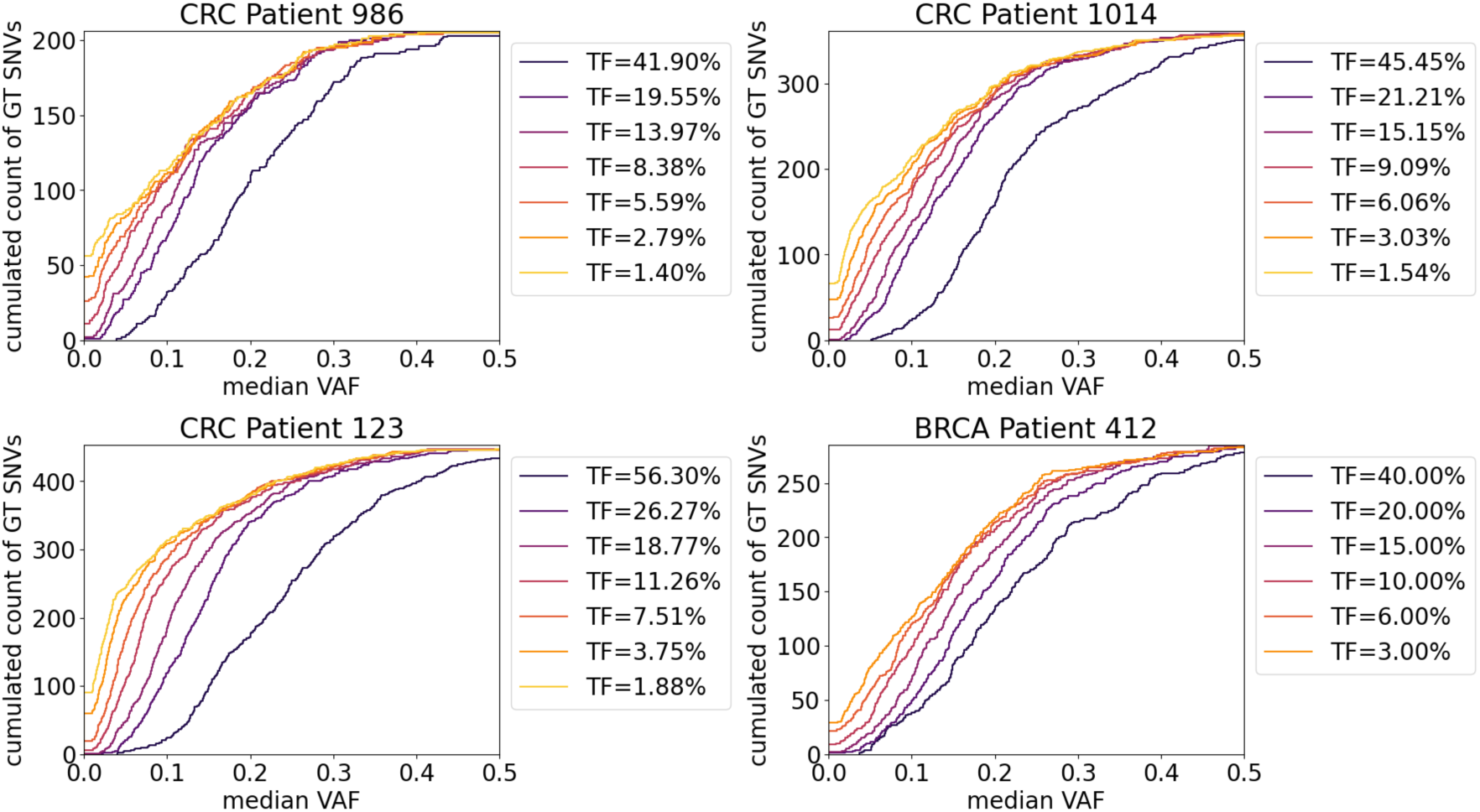
Cumulative variant allele frequency of pseudo ground truths Single Nucleotide Variants in each dilution series (A) for the four cancer patients. For each ground truth, the median variant allele frequency value reported by all nine studied callers is considered. The sample with the highest TF for each patient corresponds to the undiluted high ctDNA sample at 70x coverage depth taken as reference to build consensus ground truths. The other samples are diluted while maintaining a fixed 150x coverage depth forming the decreasing signal series (A). TF: Tumor Fraction. ctDNA: circulating tumor DNA.

**Supplementary Figure 3:**
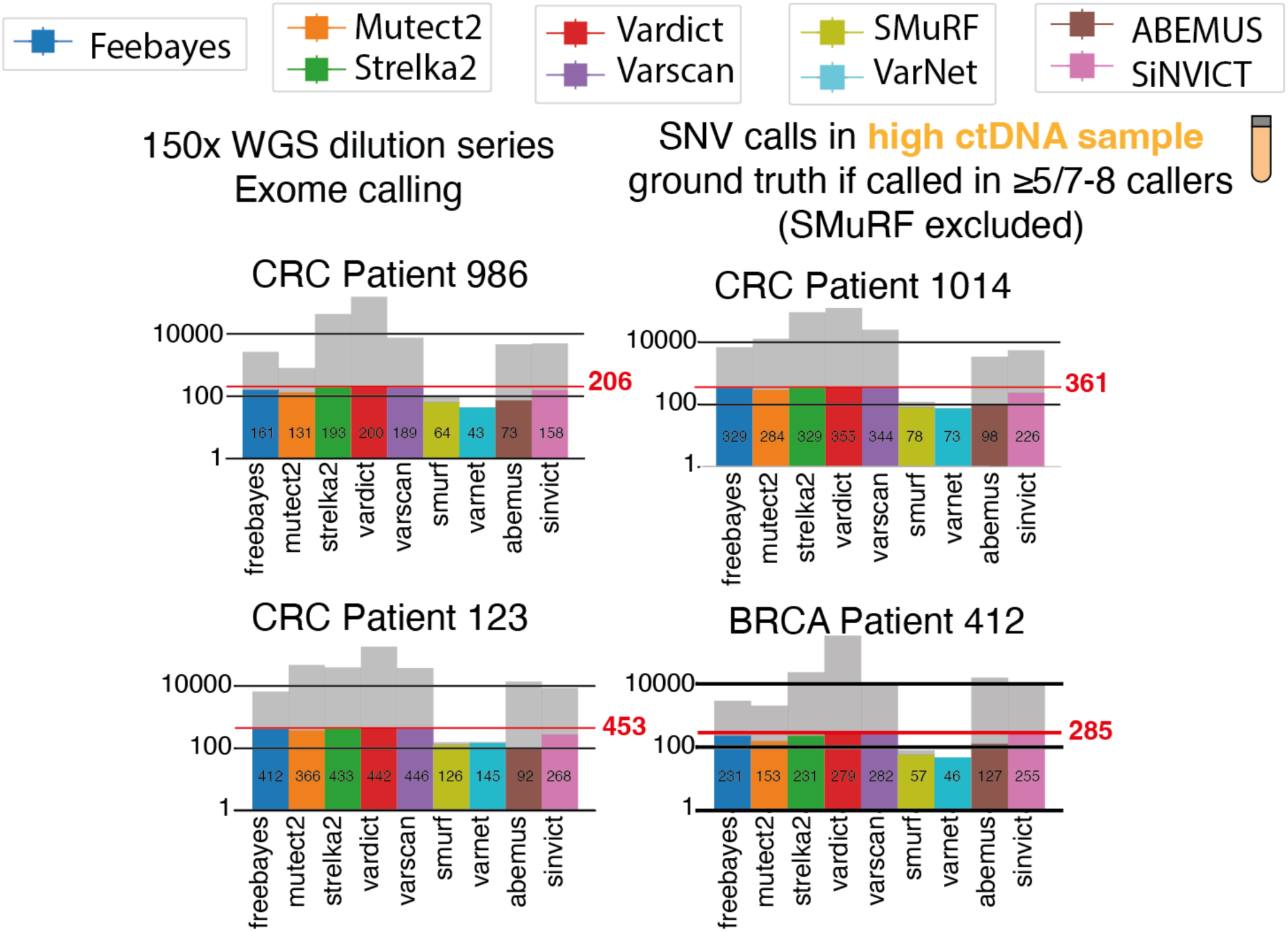
Pseudo ground truth repartition from consensus calling on the high circulating tumor DNA sample. Bar plots showing for each studied patient the number of Single Nucleotide Variants passing default filters and thresholds called in the exomic regions of the ∼100x WGS high ctDNA sample. The figure is log-scaled on the y-axis due to the wide range of calls that can be predicted by each method. Calls called by more than 5 callers among 8 callers are considered ground truths. Note that SMuRF is excluded for ground truth generation as it is an ensemble caller. Calls falling within the pseudo ground truths list are colored (True Positives) while others are shown in grey (False Positives). The red line indicates the number of consensus pseudo ground truths calls. ctDNA: circulating tumor DNA.

**Supplementary Figure 4:**
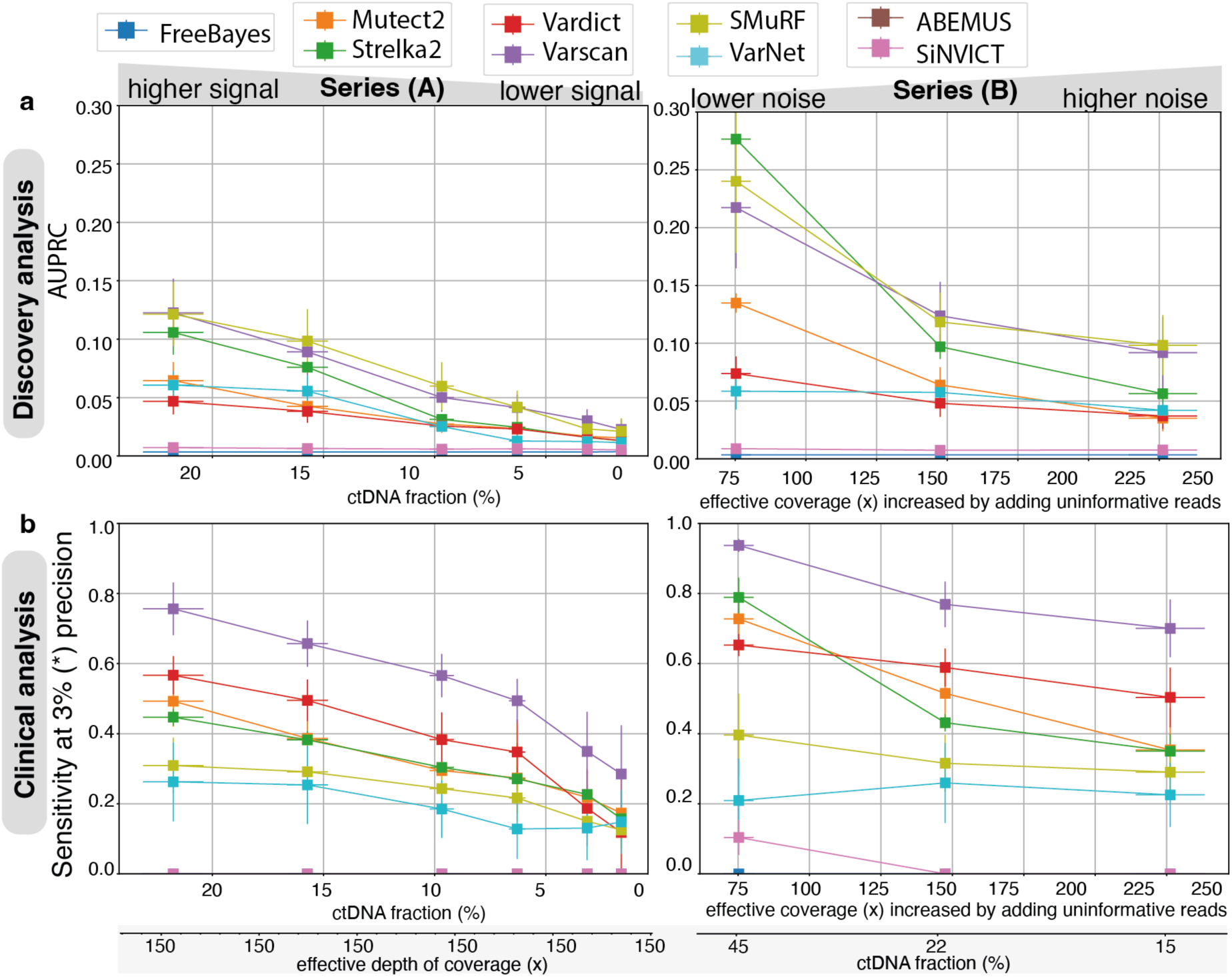
Performance results in INDEL calling on created 150x dilution series over the four studied cancer patients. Callers performance on INDELs in exonic regions averaged on the n=4 patients in both benchmark series: the decreasing signal series (A) where TF decreases at a fixed 150x depth of coverage (left, 6 samples) and the increasing noise series (B) where coverage artificially increases (from 70x to 250x) at a fixed number of informative reads (70x) (right, 3 samples). Callers are evaluated in two application settings: **(a)** discovery analysis with the Area Under Precision Recall Curve metric and **(b)** clinical analysis with the sensitivity at a precision fixed to 3%. ABEMUS is not included as it does not handle INDEL calling. The vertical errors bars show the standard error of the mean (s.e.m.) of the averaged metric curves and the horizontal error bars show the s.e.m. of the ctDNA fraction estimates among patients (n=4). INDEL: Insertion/Deletion. ctDNA: circulating tumor DNA.

**Supplementary Figure 5a:**
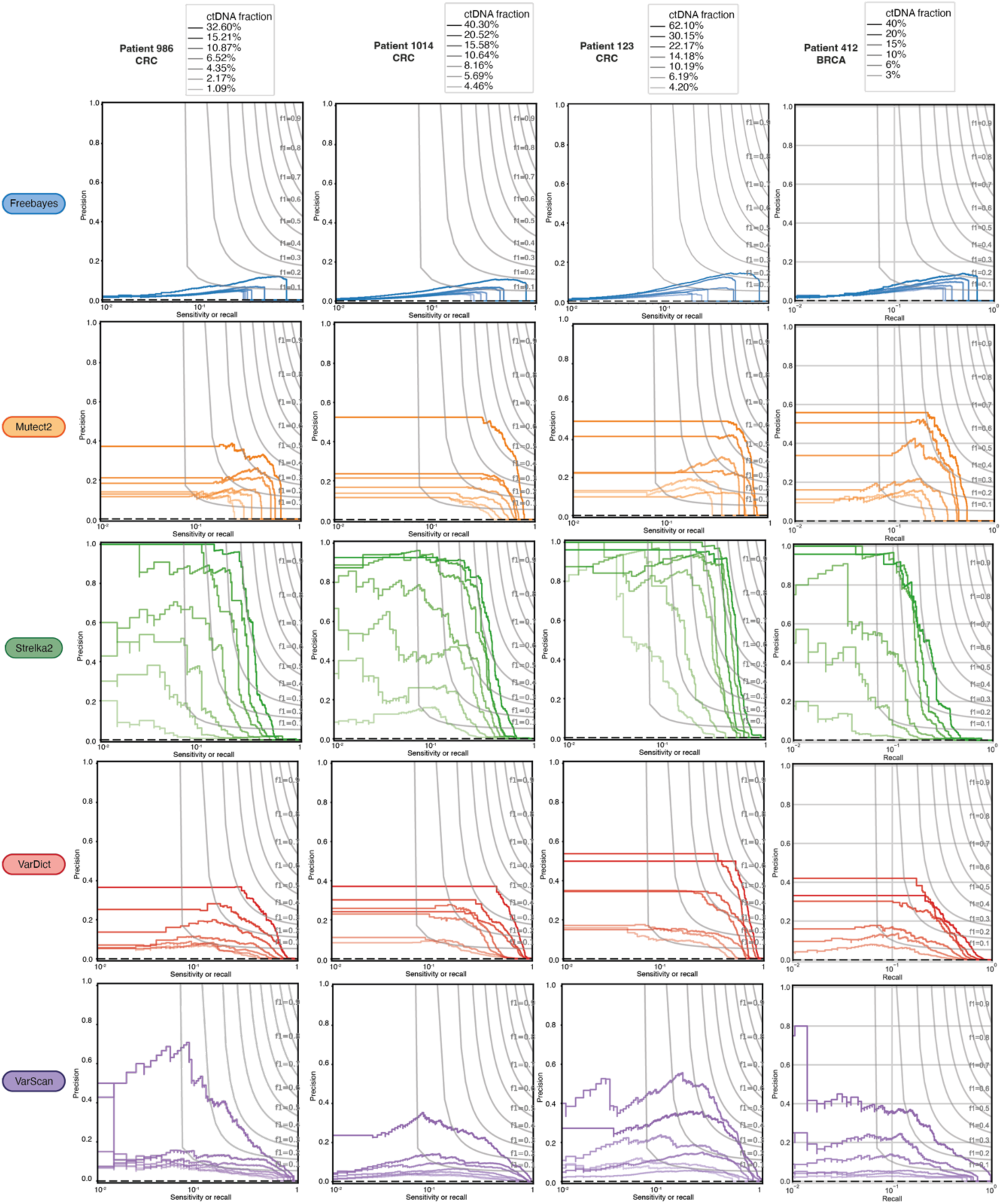
Precision-Recall curves for each somatic mutation caller for Single Nucleotide Variants on each individual studied cancer patient. Namely, the patients are: CRC patient 986, CRC patient 1014, CRC patient 123, and BRCA patient 412. Callers: FreeBayes, Mutect2, Strelka2, VarDict, VarScan. CRC: Colorectal Cancer, BRCA: Breast Cancer.

**Supplementary Figure 5b:**
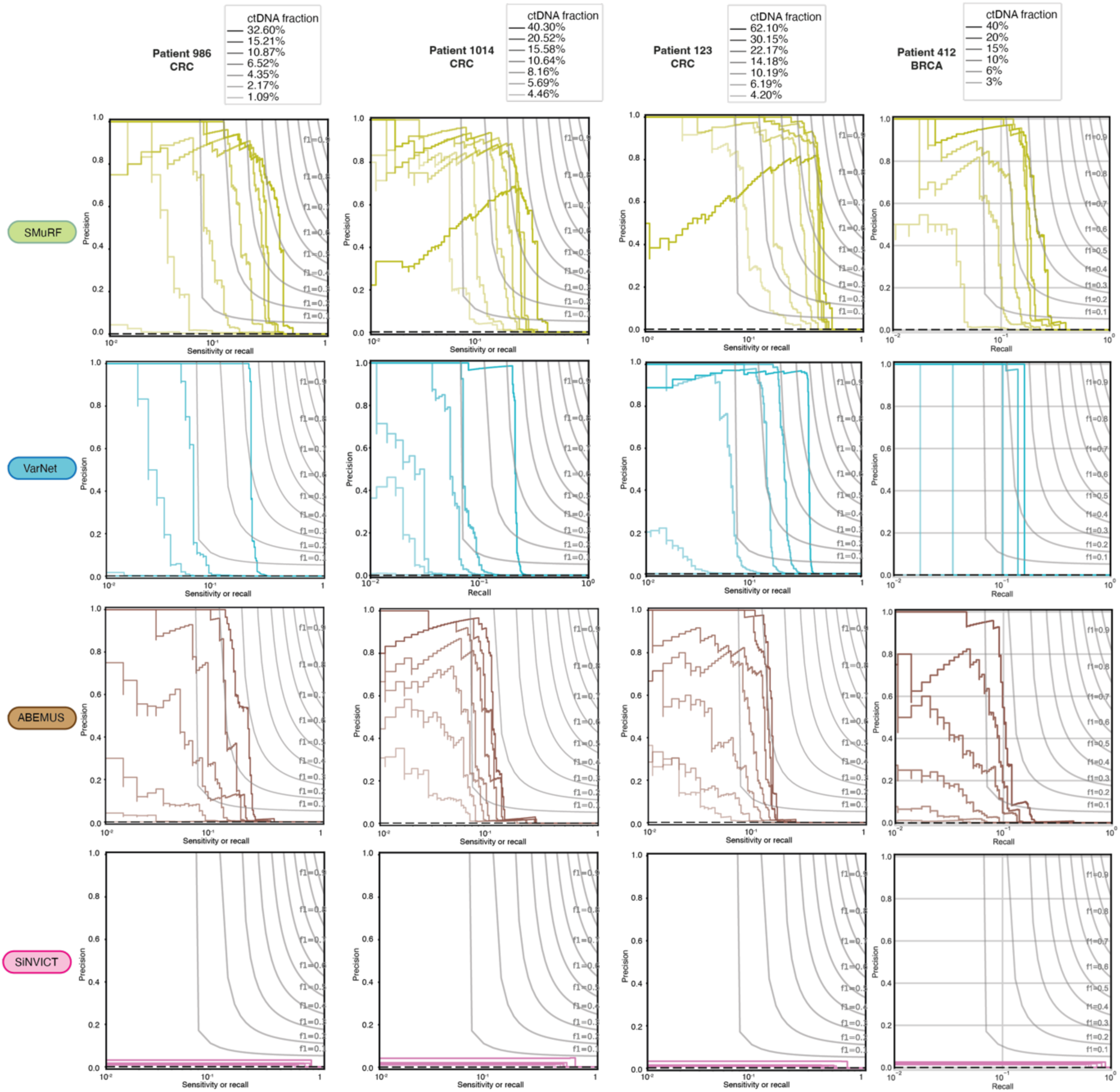
Precision-Recall curves for each somatic mutation caller for Single Nucleotide Variants on each individual studied cancer patient. Namely, the patients are: CRC patient 986, CRC patient 1014, CRC patient 123, and BRCA patient 412. Callers: SMuRF, VarNet, ABEMUS, SiNVICT. CRC: Colorectal Cancer, BRCA: Breast Cancer.

**Supplementary Figure 6:**
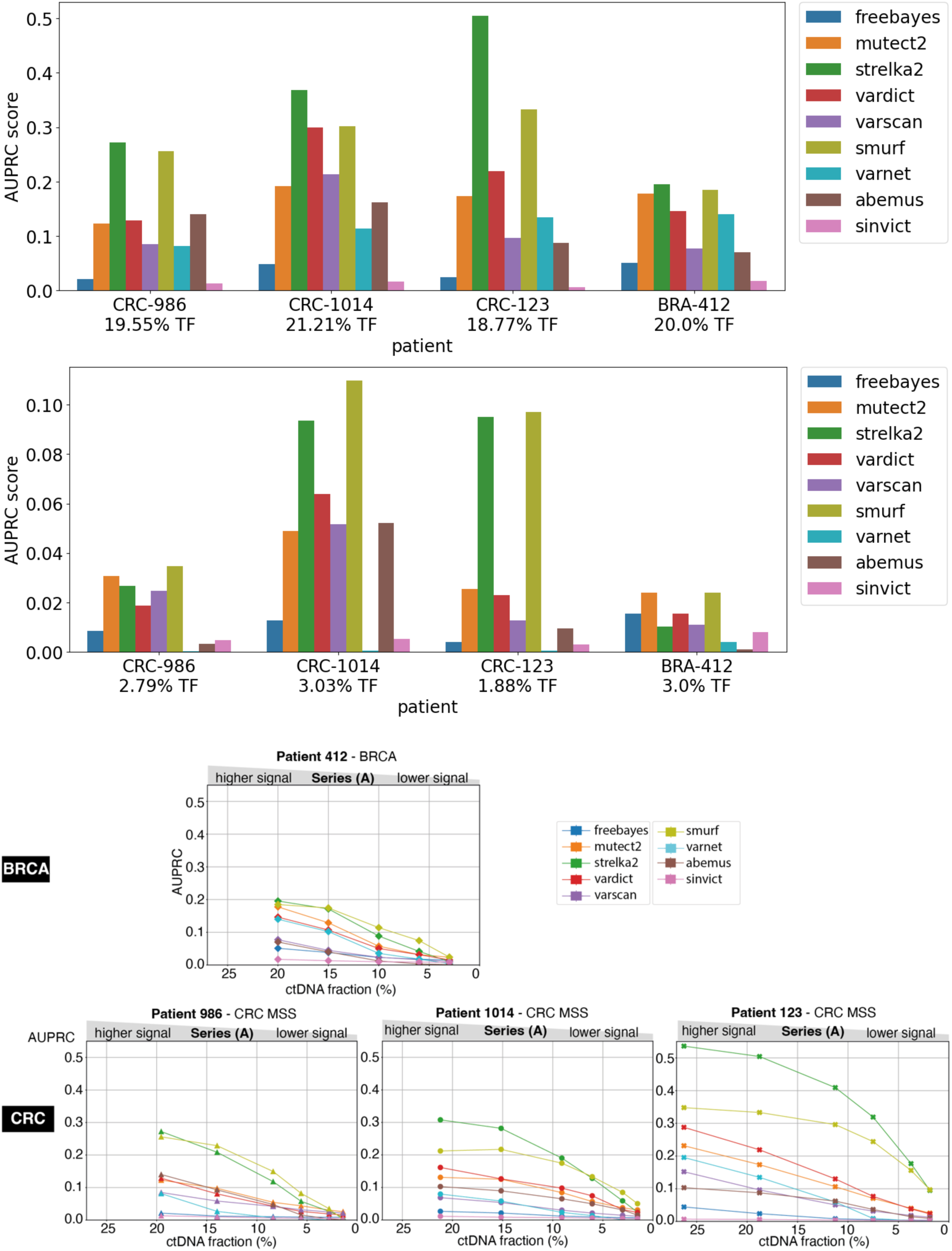
Results concordance between the two widespread Colorectal and Breast Cancer types. Callers’ performance on Single Nucleotide Variants in exonic regions for each individual patient is displayed in the decreasing signal series (A). The top barplots show the side-by-side AUPRC for each patient in one high TF (TF∼20%) and one low TF (TF∼3%) benchmark dilution. The bottom plots show the AUPRC across all benchmark dilutions for each patient. AUPRC: Area Under Precision-Recall Curve, TF: Tumor Fraction.

**Supplementary Figure 7:**
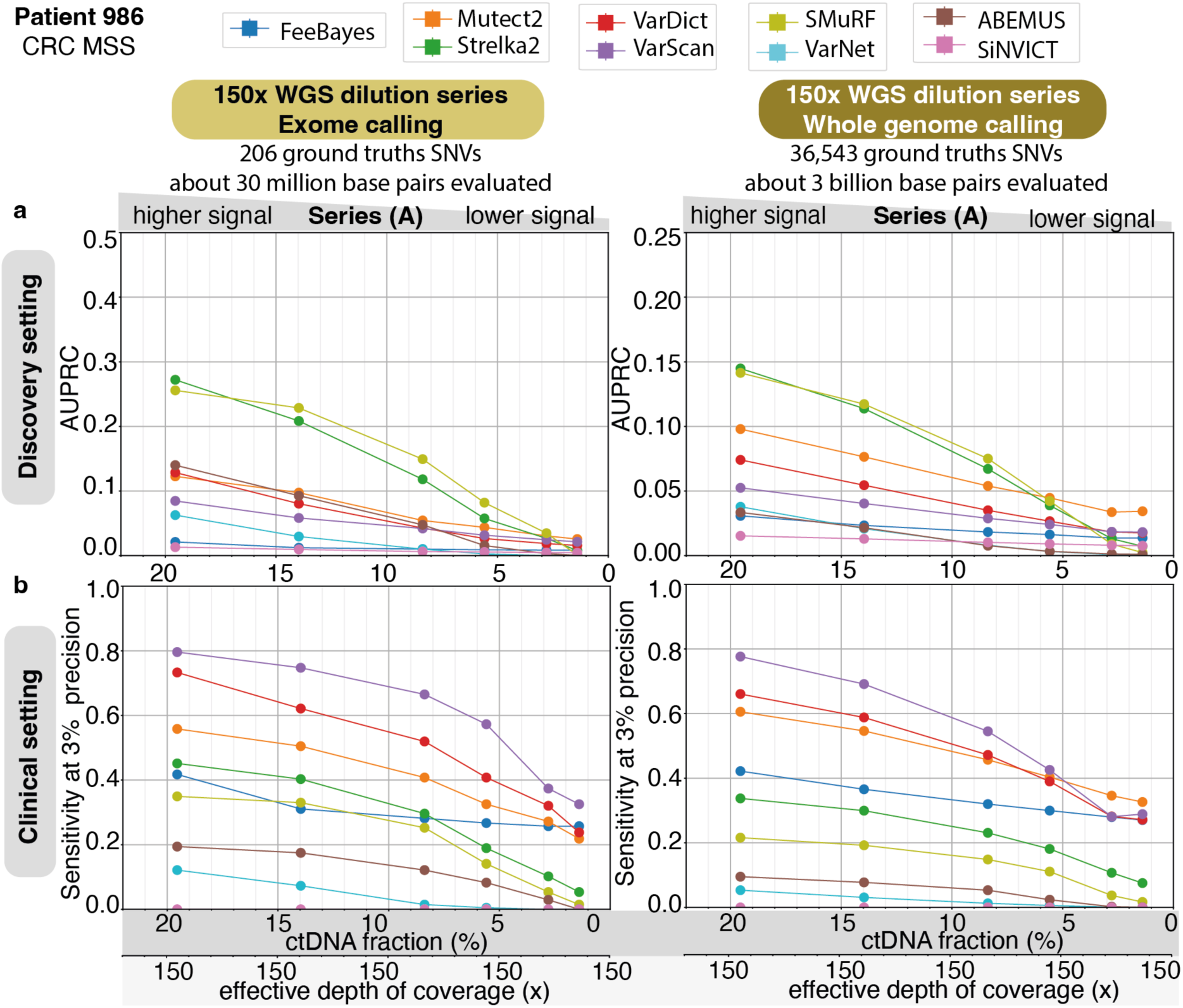
Results concordance between exome and whole genome calling. Callers’ performance on Single Nucleotide Variants in Colorectal Cancer patient 986 in the decreasing signal dilution series (A) on 150x Whole Genome Sequencing samples when applying callers either on exome (left) or on whole genome (right) in both application settings: for **(a)** discovery analysis with the Area Under Precision-Recall Curve metric and for **(b)** clinical analysis with the sensitivity at precision to 3%.

**Supplementary Figure 8:**
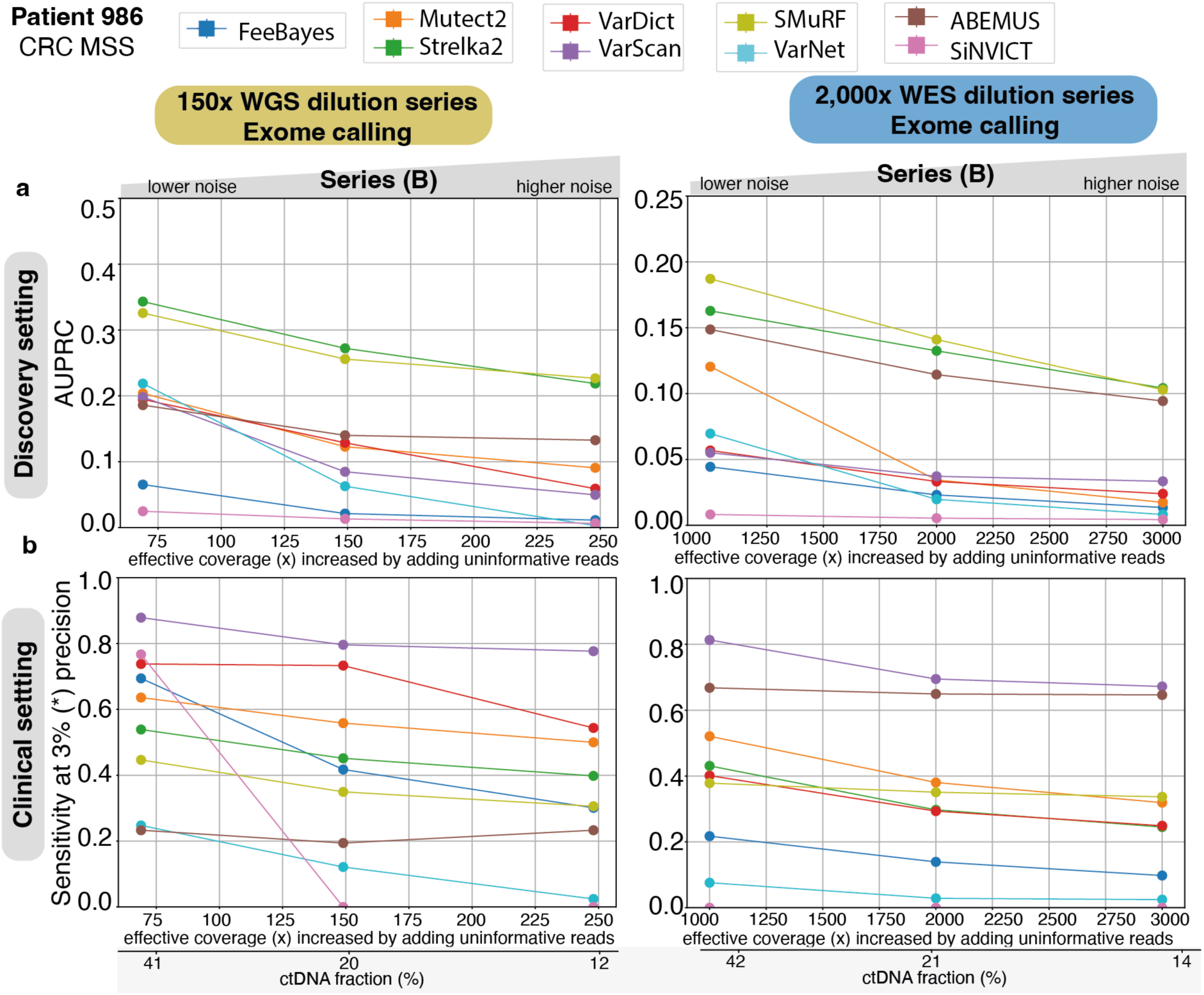
Impact of sequencing depth (150x vs 2,000x) on callers’ performance in increasing noise series (B). Callers’ performance on Single Nucleotide Variants in exonic regions of Colorectal Cancer patient 986 in dilution series (B) at two different fixed coverage levels: 150x (left, derived from Whole Genome Sequencing samples) and 2,000x (right, derived from Whole Exome Sequencing samples). Callers are evaluated in two application settings: **(a)** discovery analysis with the Area Under Precision-Recall Curve metric with a zoomed version of the ctDNA fraction range [1-5%] and **(b)** clinical analysis with the sensitivity at a precision fixed to 3%. ctDNA: circulating tumor DNA.

**Supplementary Table 1:** Tumor Fraction estimates of initial samples before dilutions. Selected cancer plasma samples detailed information to build a reliable TF estimate for each initial deep WGS sample. For high ctDNA samples, the TF is calculated as the mean of ichorCNA [42] TF and twice the median VAF of known cancer driver mutations which were first screened in the matched deep targeted sample. For ultra-low ctDNA samples, the TF is taken as the mean of the max VAF and of twice the median VAF of those known cancer driver mutations. The TF estimate relied on VAF in this case as the ichorCNA [42] TF estimate found was not consistent between the matched shallow and deep WGS samples.

| *Patient ID*<br>*Cancer type*<br>*and subtype* | *Sample characteristics*<br>*Timepoint*<br>*Sequencing type and cov* | *Tumor*<br>*Fraction*<br>*(TF)*<br>*estimate* | *Intermediate values to build TF estimate*<br>*and additional values for orthogonal*<br>*verification* |
| --- | --- | --- | --- |
| Patient 986<br>CRC MSS | **high ctDNA sample**<br>timepoint: 40 days<br>deep WGS (71.63x cov) | TF = 32.6% | ichorCNA: **23%**<br>median of 4 purity estimates: **41.9%**<br>max VAF: **40.7%**<br>2 x median VAF: 2 x 21.1% = **42.2%** |
|  | **ultra-low ctDNA sample**<br>timepoint: 454 days<br>deep WGS (~100x cov) | TF = 0% | ichorCNA: **8%**<br>ichorCNA (matched lpWGS): **0%**<br>max VAF: **0%**<br>2 x median VAF: 2 x 0% = **0%** |
| Patient 1014<br>CRC MSS | **high ctDNA sample**<br>timepoint: 901 days<br>deep WGS (80.84x cov) | TF = 40.3% | ichorCNA: **40%**<br>median of 4 purity estimates: **45.5%**<br>max VAF: **44.8%**<br>2 x median VAF: 2 x 20.3% = **40.6%** |
|  | **ultra-low ctDNA sample**<br>timepoint: 800 days<br>deep WGS (~100x cov) | TF = 2.56% | ichorCNA: **5.1%**<br>ichorCNA (matched lpWGS): **0%**<br>max VAF: **1.9%**<br>2 x median VAF: 2 x 1.61% = **3.22%** |
| Patient 123<br>CRC MSS | **high ctDNA sample**<br>timepoint: 1641 days<br>deep WGS (71.63x cov) | TF = 62.1% | ichorCNA: **65.4%**<br>median of 4 purity estimates: **47%**<br>max VAF: **56.5%**<br>2 x median VAF: 2 x 29.4% = **58.8%** |
|  | **ultra-low ctDNA sample**<br>timepoint: 1745 days<br>deep WGS (~100x cov) | TF = 3.15% | ichorCNA: **6.1%**<br>ichorCNA (matched lpWGS): **0%**<br>max VAF: **4.1%**<br>2 x median VAF: 2 x 1.10% = **2.20%** |
| Patient 412<br>BRCA | **high ctDNA sample**<br>timepoint: 1414 days<br>deep WGS (71.63x cov) | TF = 67.6% | ichorCNA: **66.4%**<br>median of 4 purity estimates: **NA**<br>max VAF: **68.9%**<br>2 x median VAF: **NA** |
|  | **ultra-low ctDNA sample**<br>timepoint: 1218 days<br>deep WGS (~100x cov) | TF = 0% | ichorCNA: **0%**<br>ichorCNA (matched lpWGS): **0%**<br>max VAF: **0%**<br>2 x median VAF: 2 x 0% = **0%** |

**Supplementary Table 2:**
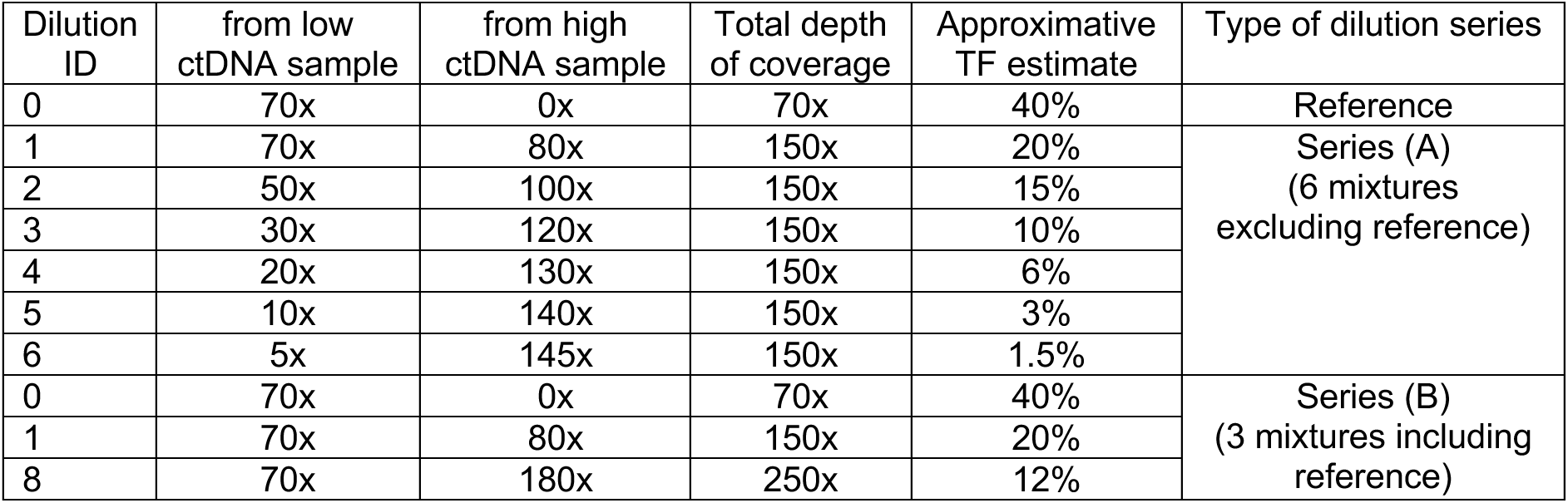
Dilution ratio choice taken in experimental design for dilution series creation at 150x whole genome sequencing. Sample ID = 0 will be the reference undiluted high ctDNA sample. From a pair of high/ultra-low ctDNA samples of one cancer patient, 8 samples are created for Series (A), Series (B), and reference sample. ctDNA: circulating tumor DNA, TF: Tumor Fraction.

**Supplementary Table 3:** Dilution ratio choice taken in experimental design for dilution series creation at 2,000x whole exome sequencing. Sample ID = 0 will be the reference undiluted high ctDNA sample. From a pair of high/ultra-low ctDNA samples of one cancer patient, 8 samples are created for Series (A), Series (B), and reference sample. ctDNA: circulating tumor DNA, TF: Tumor Fraction.

| Dilution ID | from low ctDNA sample | from high ctDNA sample | Total depth of coverage | Approximative TF estimate | Type of dilution series |
| --- | --- | --- | --- | --- | --- |
| 0 | 1,000x | 0x | 1,000x | 40% | Reference |
| 1 | 1,000x | 1,000x | 2,000x | 20% | Series (A)<br>(6 mixtures excluding reference) |
| 2 | 750x | 1,250x | 2,000x | 15% |  |
| 3 | 500x | 1,500x | 2,000x | 10% |  |
| 4 | 400x | 1,600x | 2,000x | 6% |  |
| 5 | 200x | 1,800x | 2,000x | 3% |  |
| 6 | 100x | 1,900x | 2,000x | 1.5% |  |
| 0 | 1,000x | 0x | 1,000x | 40% | Series (B)<br>(3 mixtures including reference) |
| 1 | 1,000x | 1,000x | 2,000x | 20% |  |
| 8 | 1,000x | 1,900x | 2,000x | 12% |  |

**Supplementary Table 4:** Studied callers underlying principles and implementation details. This table contains a concise description of each studied caller’s underlying principle as well as whether it handle short INDEL calling, the computing resources required to apply it on a dilution series of eight 2,000x whole exome sequencing cfDNA samples in plasma/normal mode, and includes some practical implementation notes. INDEL: Insertion/Deletion. (*) INDELs narrowed down to short INDELs ≤35bp. (**) Resources numbers given for a 2,000x WES cfDNA dilution series of 8 samples (∼100Gb each) with matched WBC sample. INDEL: Insertion/Deletion.

| *Tissue*<br>*VS*<br>*cfDNA* | *Somatic*<br>*Mutation*<br>*Caller* | *Underlying principle* | *handles*<br>*INDEL*<br>*calling*<br>*(\*)?* | *For a 2,000x WES mixture series*<br>*of 8 samples* |  |  | *Implementation notes* |
| --- | --- | --- | --- | --- | --- | --- | --- |
|  |  |  |  | *Computing*<br>*time*<br>*(\*\*)* | *Computing*<br>*resources*<br>*(\*\*)* | *Temporary*<br>*storage*<br>*(\*\*)* |  |
| **Tissue**<br>**based**<br>**methods** | **FreeBayes** | Bayesian haplotype-based caller using genotyping algorithm considering likelihood and prior probabilities. | ✓ | 1 job per sample<br>each job:<br>3,5 days<br>for 5<br>callers | 1 job per sample<br>each job:<br>8Gb RAM<br>112 CPUs | 1 job per sample<br>each job:<br>1Tb | -bash and Python package<br>-integrated in bcbio-nextgen pipeline. All 5 callers can be run at once.<br>-intermediate files use 4 times the size of analysed data |
|  | **Mutect2** | Likelihood model using local assembly, tumor-specific statistical modelling, and machine learning-based error correction. | ✓ |  |  |  |  |
|  | **Strelka2** | Mixture model that first identifies potential variants using local assembly and then applies a Bayesian classifier to differentiate true mutations from sequencing artifacts. | ✓ |  |  |  |  |
|  | **VarDict** | Heuristic variant frequency-based approach using Fisher exact test on read counts after performing local realignments. | ✓ |  |  |  |  |
|  | **VarScan** | Heuristic variant frequency-based approach using statistical tests, such as the Fisher exact test or chi-square test. It also considers read depth and base quality. | ✓ |  |  |  |  |
|  | **SMuRF** | ensemble Random Forest classifier taking the 5 previous callers outputs as input. The model is trained on the ICGC dataset [19]. | ✓ | 1 job per sample<br>each job:<br>1min | 1 job per sample<br>each job:<br>20Gb RAM<br>112 CPUs | 1 job per sample<br>each job:<br>30Mb | -R package<br>-requires to run 5 callers above beforehand |
|  | **VarNet** | Convolution Neural Network model encoding each candidate mutation site into an image of 70 bp and of 100 randomly selected tumor-normal reads. Weakly-supervised learning on TCGA with SMuRF labels. | ✓ | 1 job per sample<br>each job:<br>5 hours | 1 job per sample<br>each job:<br>18Gb RAM<br>112 CPUs | 1 job per sample<br>each job:<br>30Mb | -Python package<br>-recommended pre-processing: marking and removing duplicates<br>-no interpretable features listed in output |
| **cfDNA**<br>**based**<br>**methods** | **ABEMUS** | two-step filtering approach using both a filter on global error measure and a filter on per-base error measure. | ✗ | 1,2 day | 1Tb RAM<br>112 CPUs | 40Gb | -R package<br>-requires a Panel of Normal (PoN) of $\geq 10$ WBC samples with the same sequencing platform and depth. No default PoN provided. |
|  | **SiNVICT** | multi-step approach on the plasma sample only using local assembly, read realignment, and a statistical Poisson model under the assumption of a background mutation rate. | ✓ | without ABRA<br>17 hours | 165Gb RAM<br>112 CPUs | 100Gb | -C++ package<br>-recommended pre-processing: ABRA very demanding.<br>-no probability score but binary output with 6 different filters<br>-does not use WBC sample, tumor only. |

